# Whole-animal connectome and cell-type complement of the three-segmented *Platynereis dumerilii* larva

**DOI:** 10.1101/2020.08.21.260984

**Authors:** Csaba Verasztó, Sanja Jasek, Martin Gühmann, Réza Shahidi, Nobuo Ueda, James David Beard, Sara Mendes, Konrad Heinz, Luis Alberto Bezares-Calderón, Elizabeth Williams, Gáspár Jékely

## Abstract

Nervous systems coordinate effectors across the body during movements. We know little about the cellular-level structure of synaptic circuits for such body-wide control. Here we describe the whole-body synaptic connectome and cell-type complement of a three-segmented larva of the marine annelid *Platynereis dumerilii*. We reconstructed and annotated over 1,500 neurons and 6,500 non-neuronal cells in a whole-body serial electron microscopy dataset. The differentiated cells fall into 180 neuronal and 90 non-neuronal cell types. We analyse the modular network architecture of the entire nervous system and describe polysynaptic pathways from 428 sensory neurons to four effector systems – ciliated cells, glands, pigment cells and muscles. The complete somatic musculature and its innervation will be described in a companion paper. We also investigated intersegmental differences in cell-type complement, descending and ascending pathways, and mechanosensory and peptidergic circuits. Our work provides the basis for understanding whole-body coordination in annelids.

## Introduction

Nervous systems coordinate behaviour, physiology and development through synaptic and neuroendocrine signalling. Signalling occurs specifically between groups of cells, organised into multilayered networks with precise synaptic and neuromodulatory connectivity (Bentley et al., 2016). Mapping such synaptic and chemical networks in blocks of neural tissue is the central aim of cellular-level connectomics (Deng et al., 2019; Helmstaedter, 2013; Morgan and Lichtman, 2013; Williams et al., 2017). For synaptic networks, connectomics requires volume imaging by serial electron microscopy (serial EM) (Schlegel et al., 2017).

The comprehensive analysis of whole-body coordination of actions by synaptic circuits would benefit from the cellular-level mapping of entire nervous and effector systems. Whole-animal synaptic connectomes have so far only been described for the nematode *Caenorhabditis elegans* (Cook et al., 2019; White et al., 1986) and the tadpole larva of the ascidian *Ciona intestinalis* (Ryan et al., 2016). Circuits spanning the entire central nervous system (CNS) have also been reconstructed in the larval CNS of the fruit fly *Drosophila melanogaster* (Carreira-Rosario et al., 2018; Miroschnikow et al., 2018; Ohyama et al., 2015; Schlegel et al., 2016) and in the three-day-old larva of the annelid *Platynereis dumerilii* (Bezares-Calderón et al., 2018; Randel et al., 2015; Verasztó et al., 2017a). Recently, whole-brain connectomics has become possible in the adult fly brain (Zheng et al., 2018).

Here we report the complete synaptic connectome and cell-type complement of a three-day-old larva (nectochaete stage) of the marine annelid *Platynereis dumerilii*. This larval stage has three trunk segments, adult and larval eyes, segmental ciliary bands and a well-developed somatic musculature. The larvae show several behaviours, including visual phototaxis (Randel et al., 2014), UV avoidance (Verasztó et al., 2018), a startle response (Bezares-Calderón et al., 2018) and coordinated ciliary activity (Verasztó et al., 2017b). Three-day-old larvae do not yet have sensory palps and other sensory appendages (cirri), they do not feed and lack visceral muscles and an enteric nervous system (Brunet et al., 2016; Williams et al., 2015). Three-day-old larvae also lack associative brain centres such as mushroom bodies, which only develop several days later (Tomer et al., 2010).

In *Platynereis* larvae, it has been possible to integrate behaviour with synapse-level maps, transgenic labelling of individual neurons, activity imaging and gene knockouts (Bezares-Calderón et al., 2018; Verasztó et al., 2018, 2017a). Cellular-resolution gene expression atlases have also been developed for different larval stages. These can increasingly be integrated with single-cell transcriptomic atlases and synaptic circuit maps (Achim et al., 2015; Asadulina et al., 2012; Randel et al., 2014; Tomer et al., 2010; Williams et al., 2017). A recent study reported the registration of a gene expression atlas on a non-synaptic resolution EM volume in the six-day-old *Platynereis* larva (Vergara et al., 2020).

We previously reported synaptic connectomes for several whole-body circuits from the three-day-old larva. These include the visual, startle, ciliomotor, nuchal organ, and neurosecretory systems (Bezares-Calderón et al., 2018; Randel et al., 2015; Shahidi et al., 2015; Verasztó et al., 2018, 2017a; Williams et al., 2017). Here, we report the complete synaptic connectome and the cell-type complement of the three-day-old *Platynereis* larva. In a companion paper, we will report the desmosomal connectome and motoneuron innervation of the somatic musculature (Jasek et al.). The analyses were based on the previously reported whole-body serial transmission EM volume. The full connectome reconstruction has now allowed us to uncover several new circuits and to consider all circuits in a whole-body context. We also found several neurons that span the entire length of the larva, highlighting the strength of a whole-body dataset. These cells and their circuits allow us to generate hypotheses on how whole-body coordination may be achieved by the larval nervous system. The connectome also allowed us to address long-standing hypotheses about the origin of the segmented annelid body-plan and explore patterns of circuit evolution.

## Results

### Serial EM reconstruction of a *Platynereis* larva

We traced and annotated all cells in a previously reported serial EM dataset of a three-day-old (72 hours post fertilisation (hpf) *Platynereis* larva (Randel et al., 2015). The dataset consists of 4,845 layers of 40 nm thin sections scanned by transmission electron microscopy (TEM). The sections span the entire body of the larva. We used the collaborative annotation toolkit CATMAID for tracing and reviewing skeleton and for annotations (Saalfeld et al., 2009; Schneider-Mizell et al., 2016). To mark the position of cell somas, we tagged the centre of each nucleus in the volume. We skeletonised cells containing projections or having an elongated morphology including muscle cells, glia and neurons. In neurons, we identified and marked presynaptic sites and connected the synapses to postsynaptic partners, in order to obtain the synaptic connectome (Figure 1, Figure 1 – figure supplement 1).

**Figure 1.**
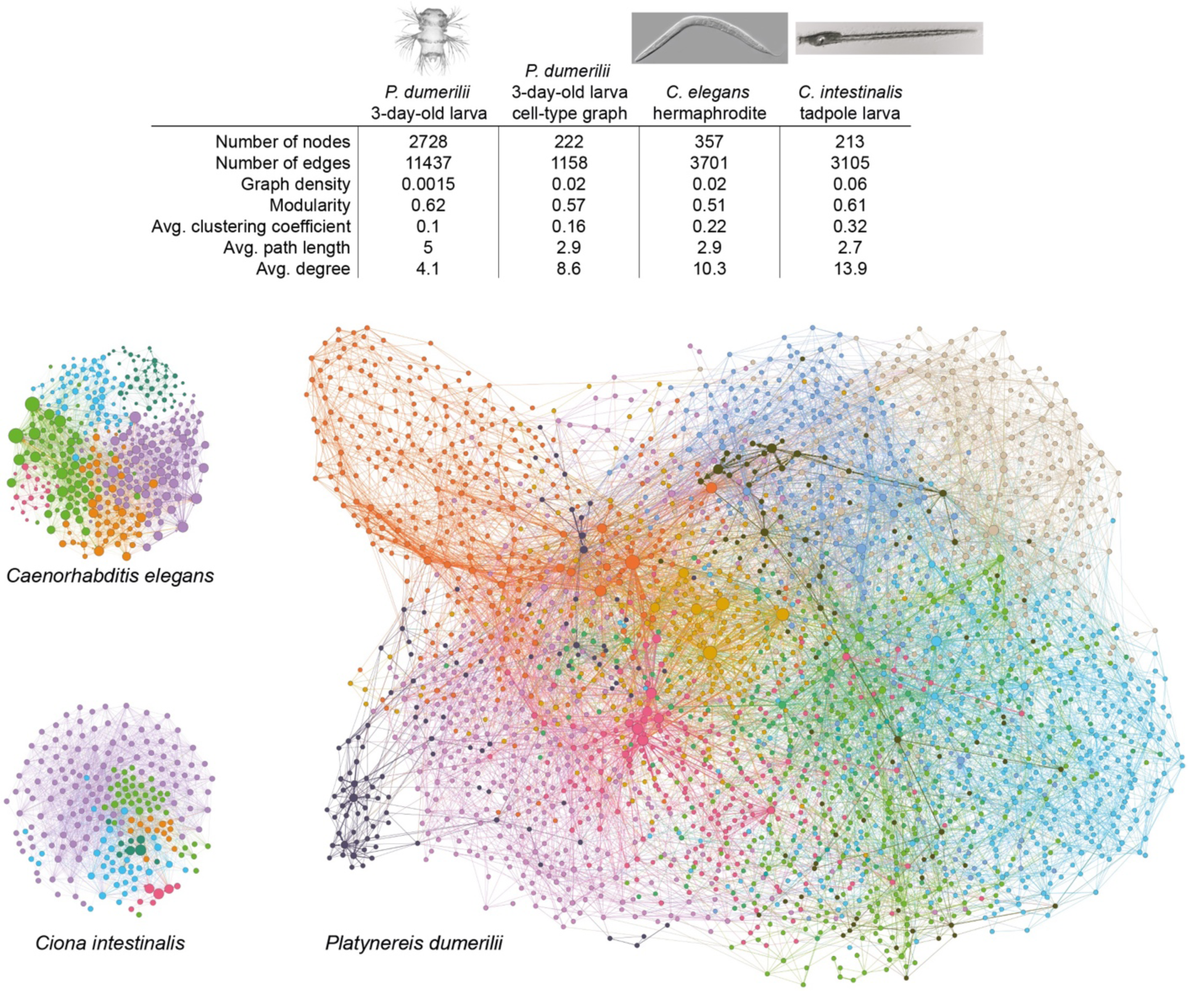
The *Platynereis dumerilii* larval connectome in comparison to the *C. intestinalis* and *C. elegans* connectomes. Graphs of the *C. elegans* hermaphrodite, *C. intestinalis* tadpole larva and *P. dumerilii* three-day-old larva chemical synapse connectomes. Nodes represent individual cells, edges represent synaptic connectivity. Arrowheads were omitted. Nodes are coloured by modules. Node sizes are proportional to weighted degree. The table shows network parameters for the connectome graphs. Figure 1 – source data 1 (adjacency matrix).

In the EM volume, we identified 8,852 cells with a soma. Their skeletons consisted of 5,519,861 nodes and had 27,943 presynaptic and 26,433 postsynaptic sites. We could not attach 16,260 fragments (861,448 nodes) to a skeleton with a soma. These fragments contained 4,070 presynaptic and 5,565 postsynaptic sites. Most of the fragments represent short skeletons of twigs that could not be traced across gaps or low-quality layers (Figure 1 – figure supplement 2). Overall, 6.4% of all nodes, 6.8% of presynaptic, and 4.7% of postsynaptic sites are on fragments and not assigned to a cell with a soma. The total construction time of all skeletons was over 3,600 hours with an additional 750 hours of review time.

### Network analysis of whole-body synaptic connectivity

For the analysis of whole-body synaptic connectivity in the larva, we selected cells with presynaptic sites – representing differentiated neurons – and their postsynaptic partners. All fragments without a soma were removed. The final synaptic connectome contains 2,728 cells connected by 11,437 edges (25,509 synapses)(Figure 1, Figure 1 – Figure Supplement 3).

The connectome is a sparsely-connected network with a graph density of 0.0015. In comparison, the *Caenorhabditis elegans* hermaphrodite connectome has a graph density of 0.02 (Cook et al., 2019) and the *Ciona intestinalis* larval connectome 0.06 (Ryan et al., 2016)(Figures 1 and 2). We also defined a grouped cell-type connectome containing only cells that were assigned to a neuronal cell type and the partners of these cell-type groups (Figure 3 and see below).

**Figure 2.**
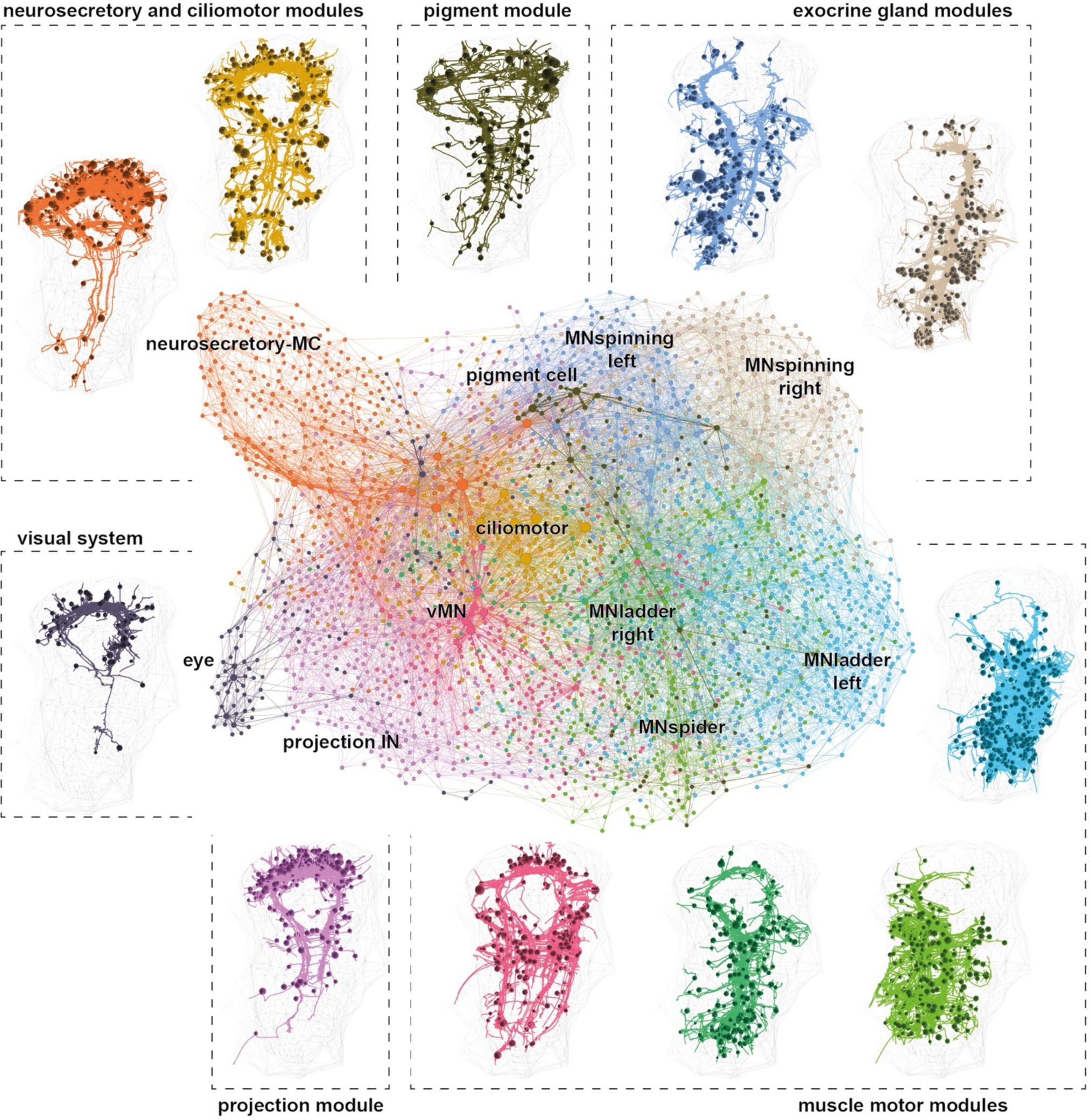
Modularity of the *Platynereis* three-day-old larval connectome. Graph representation of the connectome coloured by modules. Nodes represent single cells, edges represent synaptic connectivity. Arrowheads are not shown for clarity. The reconstructed cells for each module are shown. Spheres represent soma positions (centred on the nuclei).

**Figure 3.**
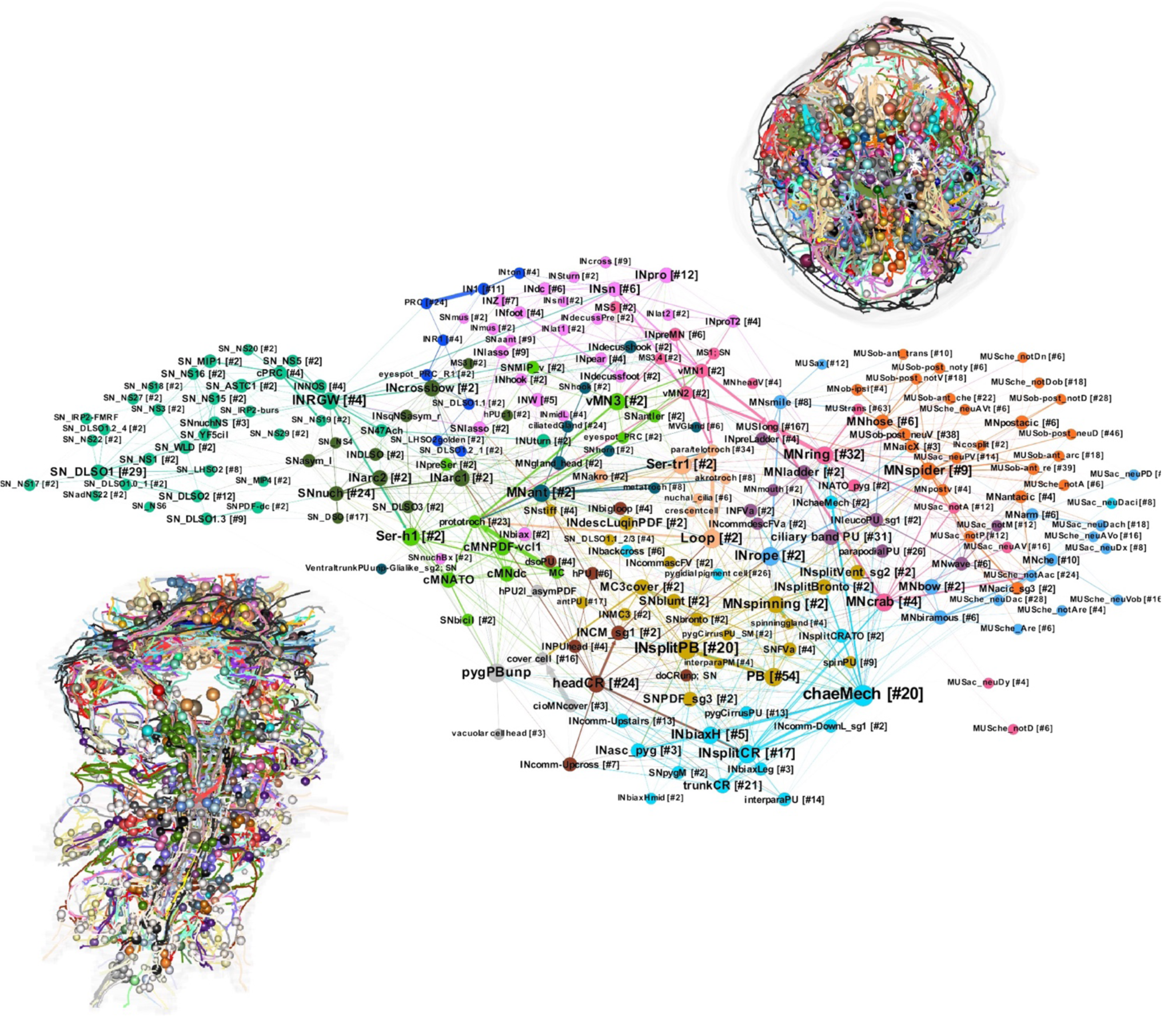
The grouped connectome of cell types. Grouped connectivity graph of cell types. Nodes represent groups of neurons, with the number of cells in square brackets. Edges represent synaptic connectivity. The width of arrows is proportional to the square root of the total number of synapses. Reconstructions of all cell types, individually coloured are shown in ventral (lower left) and anterior (upper right) view. Figure 3 – source data 1. Gephi network file of the grouped cell-type connectome.

Next we used community detection to delineate more strongly connected subgraphs within the connectome. This analysis combined with force-field-based clustering revealed several modules that may represent functional units (Figures 1 and 2). We named the modules based on their primary effector organs or other dominant anatomical characters. The modules recovered include the anterior neurosecretory centre with head ciliomotor neurons, the trunk ciliomotor system, the visual circuit, a module innervating head pigment cells, a left and right module for the innervation of segmental exocrine glands, a module of projection interneurons and four muscle-motor modules. It should be noted that community analysis can produce different numbers of modules by merging or further subdivisions, depending on the parameters used.

To identify nodes of potential functional importance we analysed node centrality in the connectome graph. We ranked nodes based on degree (number of pre- and postsynaptic partners), page rank (a measure of the number and importance of incoming links) and other measures (Figure 1 – figure supplement 2, Supplementary Table 1). We also ranked the edges connecting nodes in the graph (Supplementary Table 1). Nodes with the highest degree included ciliomotor neurons (e.g. Loop, Ser-tr1)(Bezares-Calderón et al., 2018; Verasztó et al., 2017a), the sensory-motoneuron pygPBunp (Verasztó et al., 2017a), and the motoneurons of exocrine glands (MNspinning, see below). Some of the strongest edges (highest number of synapses) were between the pigment-cell motoneuron cioMNcover and prototroch pigment cells, MNspinning and exocrine glands, and the MC cell and ciliated cells (Supplementary Table 1).

### Classification of cell types

To classify the reconstructed cells into neuronal and other cell types, we used various approaches. For a morphological classification of neurons based on the similarity of their skeleton arbots, we used Sholl analysis (Sholl, 1953) and NBLAST as implemented in CATMAID (Schneider-Mizell et al., 2016) and the `natverse` (Bates et al., 2020; Costa et al., 2016), respectively. Hierarchical clustering of distance matrices obtained from NBLAST and Sholl analysis often delineated neurons with similar arbors that we consider cell types (shown for the motoneurons in Figure 3 – figure supplement 1). These analyses however, do not use ultrastructural details (e.g. presence of sensory cilia), which we consider important for classifying cell-types. Likewise, clustering based on connectivity alone did not give sufficient resolution (data not shown).

We therefore manually classified neurons into cell types and derived a grouped cell-type graph (Figure 3). Our classification considers both morphology and connectivity. We used a combination of five criteria: i) the position of neuron somata, ii) the morphology of axon projections (e.g. branching pattern, decussation, ascending or descending), iii) the ultrastructure of sensory specialisations (e.g. number and type of cilia, microvilli – for sensory neurons only), iv) neuropeptide content as determined by the siGOLD immunolabelling method (Shahidi et al., 2015), and v) synaptic connectivity. We also required left-right symmetry for a group of similar neurons to classify as a cell type except for a few clearly asymmetric neurons (e.g. SN_YF5cil, pygPBunp). Based on these criteria, we classified 892 neurons into 180 cell types (Table 1, Video 1, Figure 3, Figure 3 – figure supplements 2 and 3).

We also categorised the remaining 7,960 cells in the larval body. 2,905 of these were classified into 90 non-neuronal cell types (Table 2). These included epithelial cells (974 cells) various pigment cells (7 types), muscle cells (852 cells of 53 types, described in the companion paper, Jasek et al.), locomotor ciliated cells (80 cells of 6 types), glial cells (3 types), gland cells (6 types), various support or sheet cells, nephridia, putative migratory cells, and parapodial cells producing or ensheathing chitin bristles (chaetae and aciculae) (Table 2, Video 2). In addition, we defined 18 broader neuronal cell groups, containing neurons of similar morphology (e.g. head decussating neurons) but in either differentiated or immature state (e.g. immature palp sensory neurons). The annotations are hierarchical, and cell groups can contain one or more differentiated cell types (Table 3).

The remaining 5,230 cells are either dividing cells (62 cells), undifferentiated cells that putatively belong to the neuronal lineage (1,692 cells) and various weakly connected or developing neurons with projections (3,476). These cells were not classified into cell types. The developing antennae contain 126 cells of which the majority (88) have few synapses and/or immature sensory dendrites. In the developing palps, we found 67 immature sensory neurons. The developing mouth (619 cells) is lined with 52 immature stomodeal sensory neurons. A further 146 immature sensory neurons and 52 non-sensory neurons occur in the dorsal head and the ventral nerve cord (VNC) (Figure 3 – figure supplement 4).

All cells were annotated in CATMAID with information representing the above categories. We also annotated all cells based on their soma position in the body (e.g. left or right side, segment 0-3, germ layer etc.; see Methods). These annotations were used to query the data and visualise subsets of cells in CATMAID or the `natverse`.

The cell-type classification allowed us to analyse a grouped synaptic connectivity graph where cells of the same type were collapsed into one node (Figure 3). This cell-type connectome included 81 sensory, 61 interneurons, 35 motoneuron types, 5 ciliated cell types, 33 muscle types, 3 glands, and 2 pigmented cell types.

By community analysis, we detected modules similar to the analysis of the full connectome graph, including ciliomotor, gland-motor, pigment-motor and muscle-motor modules (Figure 3). This cell-type graph has network parameters very similar to the *C. elegans* and *C. intestinalis* connectome graphs (Figure 1), suggesting a similar overall circuit organisation at the level of cell types, but with an order of magnitude more cells in *Platynereis*.

### Neurotransmitter and neuropeptide identities

Through a combination of *in situ* hybridisations, transgenic labelling and serial multiplex immunogold (siGOLD) labelling done previously (Bezares-Calderón et al., 2018; Conzelmann et al., 2013; Jékely et al., 2008; Randel et al., 2014; Shahidi et al., 2015; Verasztó et al., 2017a; Vergara et al., 2017), we assigned neurotransmitters or neuromodulators to several identified neurons. These transmitters and modulators could now be mapped to the whole connectome (Figure 4). We annotated 18 cholinergic, 4 serotonergic, 1 dopaminergic, 1 adrenergic, 69 glutamatergic neurons and 123 neurons expressing one of 13 different neuropeptides (pigment dispersing factor – 31, allatotropin/orexin – 23, leucokinin – 4, proenkephalin – 2, FVamide – 15, FMRFamide – 11, other RF/RYamide – 11, myoinhibitory peptide – 6, achatin – 1, RGWamide – 6, MLD/pedal peptide – 4, IRP2 – 2, WLD – 2, FVRIamide – 5 neurons). Neuropeptides occur in sensory, motor and interneurons.

**Figure 4.**
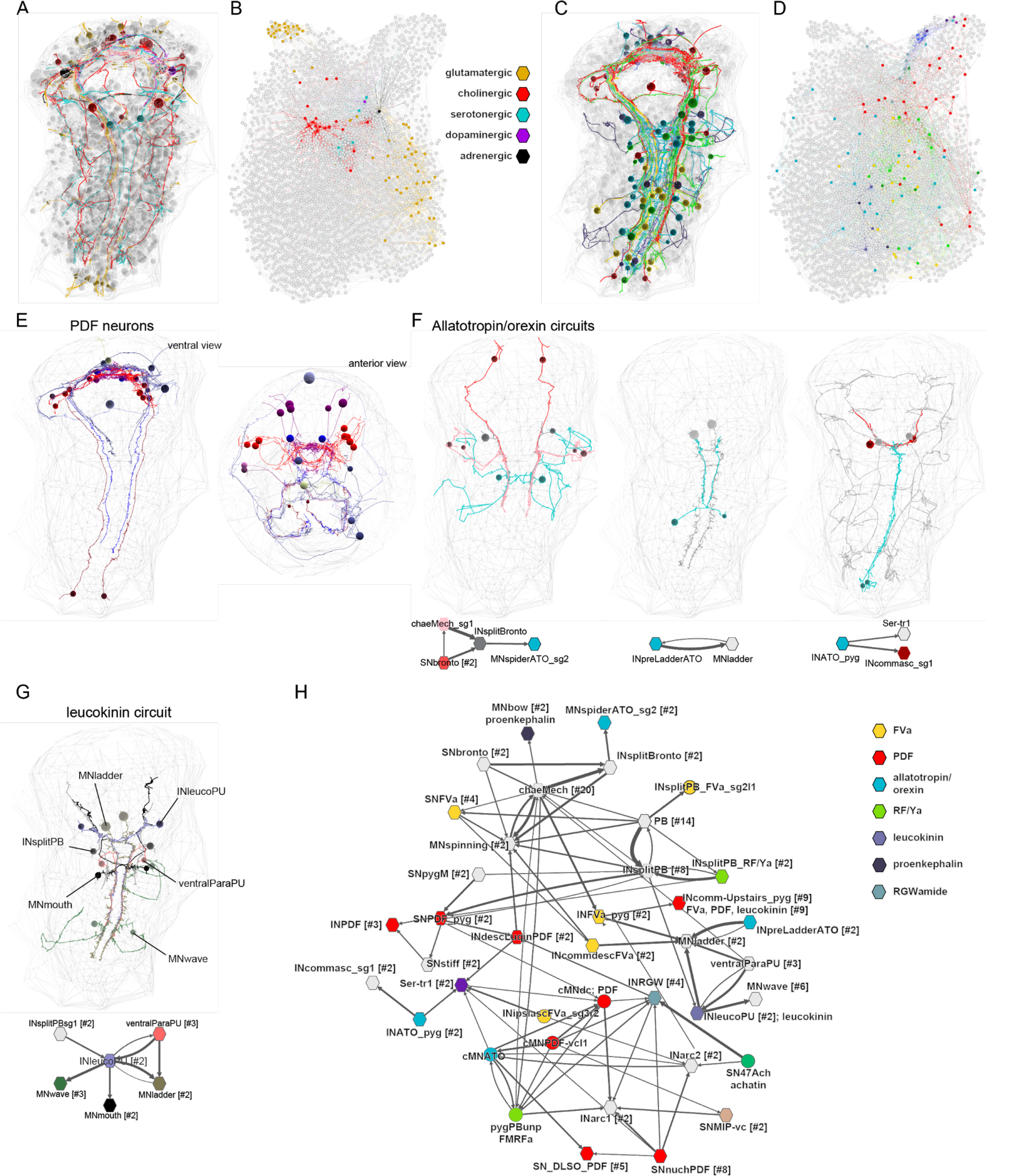
Mapping of neurotransmitters and neuropeptides to the connectome. (A) Reconstructed neurons (in colour) with a known neurotransmitter profile. (B) Neurons with a known neurotransmitter profile mapped to the connectome graph. (C) Reconstructed neurons (in colour) with a known neuropeptide profile. (D) Neurons with a known neuropeptide profile mapped to the connectome graph. (E) Neurons expressing pigment dispersing factor (PDF), ventral and anterior views. (F) Local circuits of cells expressing the allatotropin/orexin neuropeptide. (G) Local circuit of two INleucoPU neurons expressing the leucokinin neuropeptide. (H) Grouped graph of neuropeptide-expressing neurons and their pre- and postsynaptic partners. All neurons with known neuropeptide expression are shown in colour.

We highlight cellular and sensory-effector circuit examples for PDF, leucokinin and allatotropin/orexin neuropeptides (Figure 4F). For example, two leucokinin-expressing interneurons (INleucoPU) in the first trunk segment are postsynaptic to mechanosensory PU cells and INsplitPB neurons and synapse on MNladder, MNmouth, and MNwave motoneurons (Figure 4G). We also defined a larger network containing several peptidergic neurons. (Figure 4H) The specific neuropeptide-expressing cells and their mini-circuits pinpoint potential sites of peptidergic modulation within the connectome.

### Multisensory convergence and interneuron-level integration

The neuronal cell types included 45 sensory neuron types with postsynaptic partners. These collectively connect to 52 primary interneuron types, defined as interneurons directly postsynaptic to sensory neurons (Figure 5). Some sensory neurons directly connect to muscle-motor, ciliomotor or pigment-motor neurons (e.g. eyespotPRC_R3, MS cells, SNhook, SNblunt) (Figure 5E, F) or to ciliated, pigmented or muscle effector cells (e.g. eyespotPRC_R3, pygPBunp, hPU2l_asymPDF, pygCirrusPU_SM)(Figure 3 – figure supplement 2). These represent more direct sensory-motor pathways.

**Figure 5.**
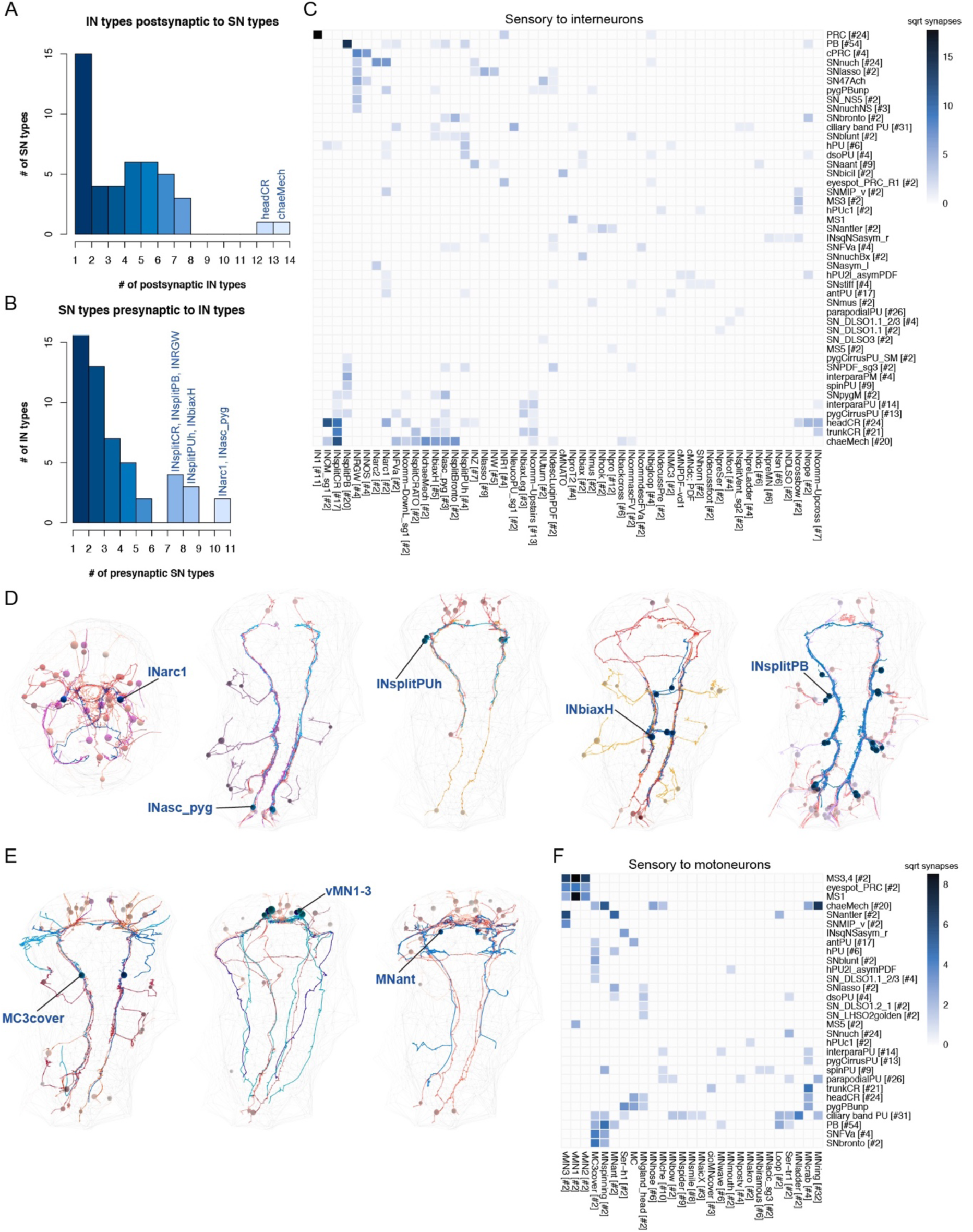
Connectivity of sensory neurons to primary interneurons and motoneurons. (A) Histogram of the number of interneurons postsynaptic to each sensory neuron type (only those with postsynaptic partners). (B) Histogram of the number of sensory neurons presynaptic to each primary interneuron type (only those with presynaptic SN partners). (C) Grouped connectivity matrix of sensory neurons and their direct postsynaptic interneuron partners. Cells are grouped by type, the number of cells in each group is indicated in square brackets. (D) EM reconstructions showing five different interneuron types with their diverse presynaptic sensory cells. (E) EM reconstructions showing three different motoneuron types with their diverse presynaptic sensory cells. (F) Grouped connectivity matrix of sensory neurons and their direct postsynaptic motoneuron partners. Cells are grouped by type, the number of cells in each group is indicated in square brackets.

The distribution of the number of partners is skewed with many sensory neurons only synapsing on one interneuron type and many interneurons only receiving input from one sensory neuron type (Figure 5A and B). At the other end of the distribution, the head CR and chaeMech sensory neurons have >10 interneuron partners, suggesting that they can recruit a more extended downstream circuit (Figure 5A).

Some interneurons and motoneurons are directly postsynaptic to several distinct sensory neuron types (up to 10)(Figure 5B-F). Such convergence suggests that these neurons can integrate multisensory inputs. Among the interneurons, INarc1, INasc_pyg, INbiaxH and INsplitPUh have >6 presynaptic sensory neuron partners (Figure 5B-D). The ventral head motoneurons (vMN) receive direct input from four sensory neuron types (MS mechanosensory, eyespotPRC_R3, SNantlerPDF, and SNMIP-vc sensory-neurosecretory neurons (Conzelmann et al., 2013))(Figure 5B, D). The MNant ciliomotor neurons are postsynaptic to three sensory neuron types in the ventral head (SNantlerPDF, SNhook, and SNhorn neurons).

The interneurons with the highest number of interneuron partners were a group of INW cells (Figure 5C). These neurons belong to the head ‘projection neurons’ module and connect to many pre- and postsynaptic interneuron types (to some only with few synapses). The INW neurons have decussating axons that delineate a V-shaped brain neuropil (not shown). This region may represent a developing integrative brain centre.

### Descending and ascending pathways connecting the brain and the VNC

Next we analysed synaptic connectivity between the brain and the ventral nerve cord. We identified all brain neurons with descending projections into the ventral nerve cord (VNC) and all trunk neurons with ascending projections into the brain. The brain has 146 such neurons, including ipsilaterally descending sensory and interneurons and decussating inter- and motoneurons. In the trunk, we found 76 ascending neurons with axons reaching the brain neuropil. The ascending and descending neurons are highly interconnected between each other and with other head and trunk neurons (Figure 6).

**Figure 6.**
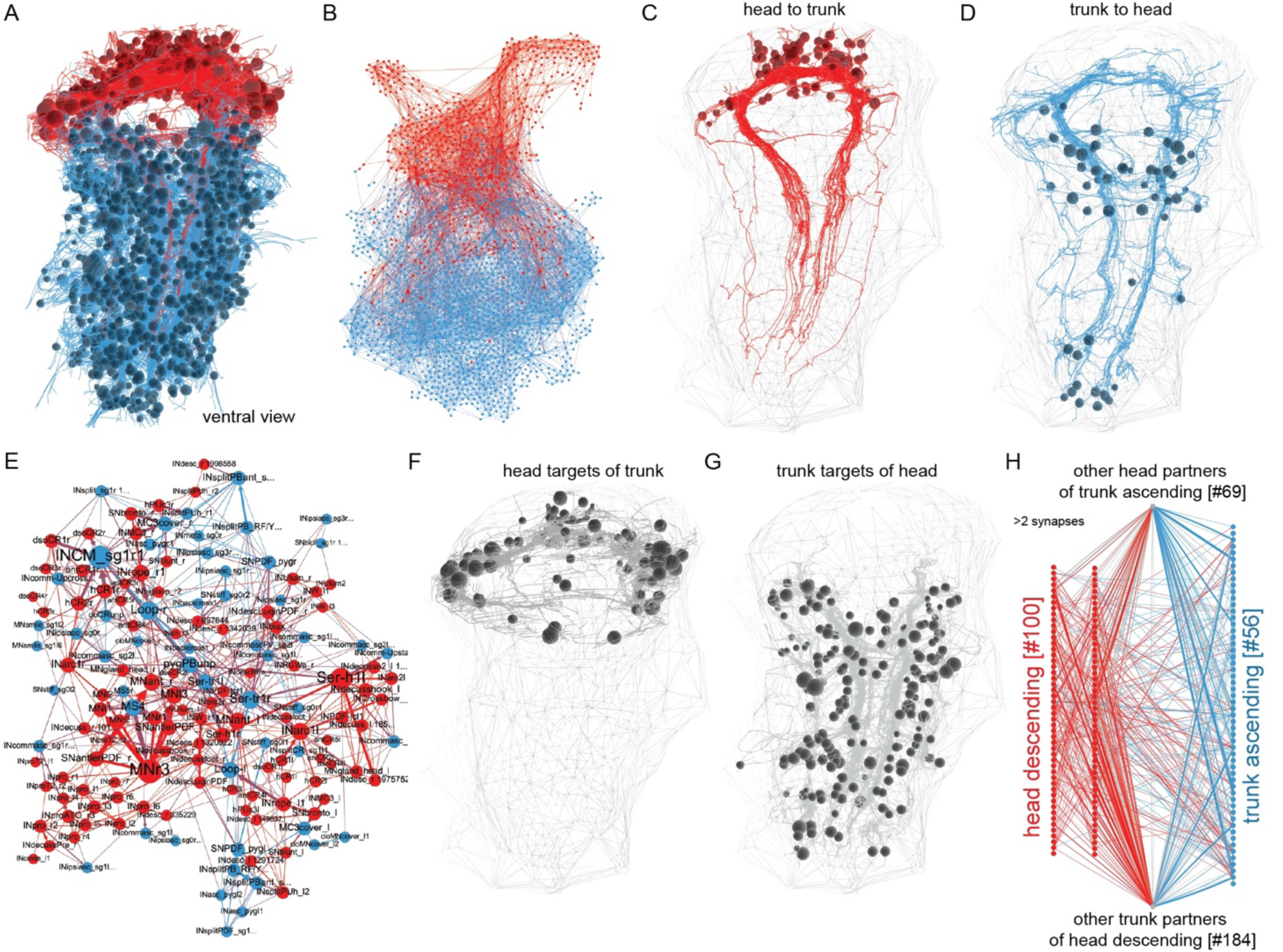
Interconnectedness of head and trunk circuits. (A) Reconstructed head (red) and trunk (blue) neurons, which are part of the connectome. (B) Graph representation of head (red) and trunk (blue) neuron connectivity. (C) Skeletons of all head neurons with descending projections into the ventral nerve cord. (D) Skeletons of all trunk neurons with ascending projections into the head. (E) Interconnectivity of head descending and trunk ascending neurons. The force-field-based layout shows how head and trunk neurons intermingle (F) All head neurons (other than in C) with synaptic contacts to trunk ascending neurons (shown in D). (G) All trunk neurons (other than in D) with synaptic contacts to head descending neurons (shown in C). (H) Network representation of synaptic connectivity of the neuron groups shown in C,D,F,G. Edges represent synaptic connections and are coloured as their source node in (H) or with a colour mixing the source and target colour in B and E (edges connecting head and trunk neurons are purple).

Some neurons connecting the trunk and the head have a global reach, with projections spanning all trunk segments and the brain (Figure 7 C-E). We identified 40 such neurons (annotation: “global_reach”), 20 of which are part of the mechanosensory girdle (see below). In the trunk, these cells only occur in the first segment and the pygidium. The pygidial neurons (e.g. cioMNcover, SNPDF_pyg, INasc_pyg, pygPBunp) span the entire VNC and terminate in the head and synapse on head neurons or on the prototroch ciliary band and its cover cells. Similarly, some head neurons have descending projections that span all trunk segments (e.g. vMN, Ser-h1, INrope, MNgland_head). The first segment contains many unique cell types with global reach (e.g. Ser-tr1, MC3cover, Loop, INsplitPBant).

**Figure 7.**
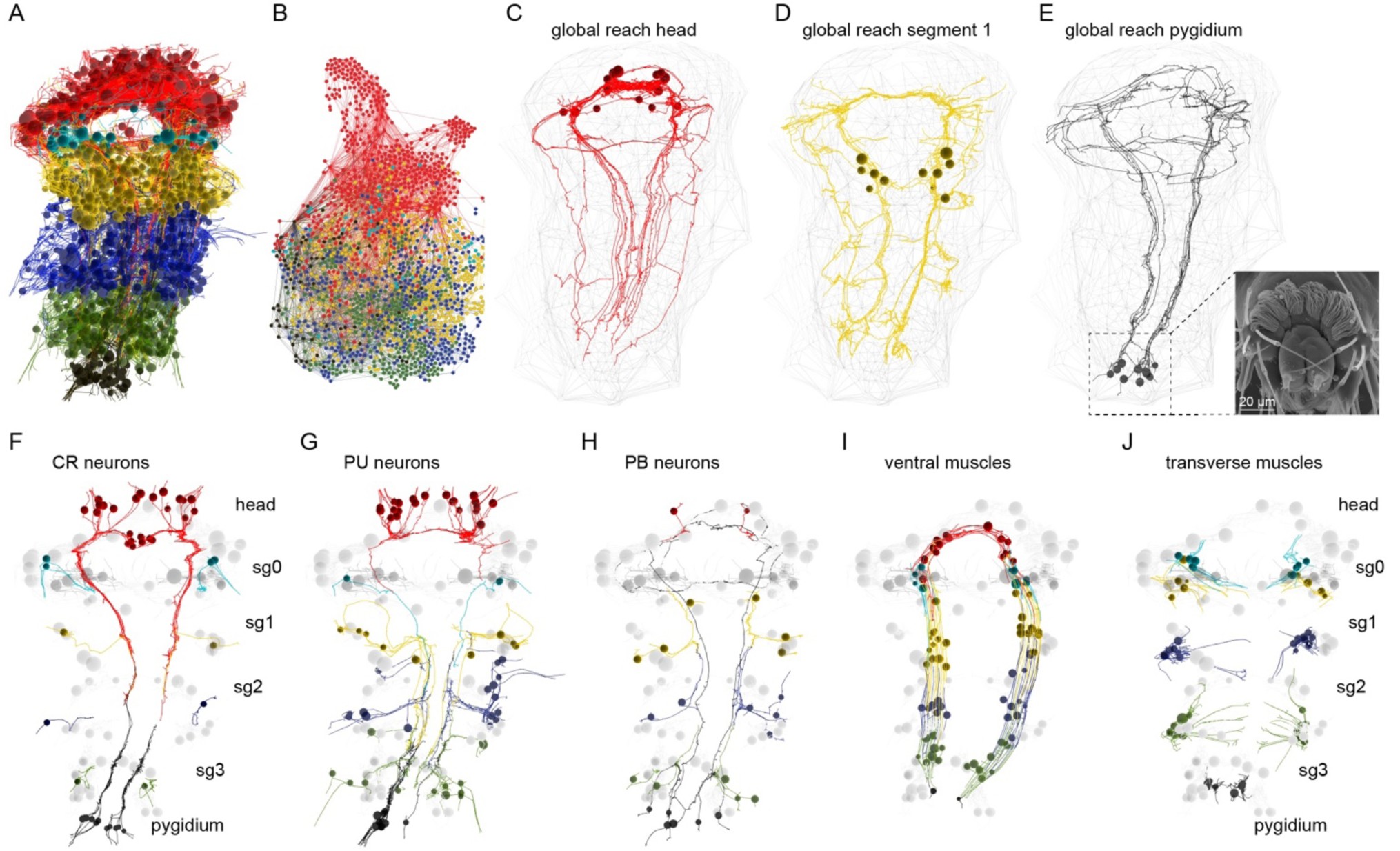
Segment-specific and segmentally iterated cell types. (A) Reconstructed cells of the connectome coloured by segment. (B) Connectome graph coloured by segment (colours same as in panel A) (C) Head-specific neurons with a global reach (D) Neurons in the first segment with a global reach. Inset: SEM image of the pygidium. (E) Neurons in the pygidium with a global reach. (F) Segmentally iterated collar receptor (CR) mechanosensory neurons. (G) Segmentally iterated uniciliated penetrating mechanosensory neurons (H) Segmentally iterated biciliated penetrating mechanosensory neurons. (I) Segmentally iterated ventral longitudinal muscles (MUSlong_V). (J) Segmentally iterated transverse muscles (MUStrans).

A small group of neurons with global reach are involved in body-wide ciliary coordination (Loop, Ser-tr1, Ser-h1)(Verasztó et al., 2017a) or coordinated intersegmental movements (INrope, vMN)(Bezares-Calderón et al., 2018; Randel et al., 2014). The function of the other descending and ascending cell types is not known and may relate to intersegmental motor coordination. Similar descending neurons coordinate trunk movements in *Drosophila* and leech. For example, the MDN descending neurons in the fly activate backward locomotion and suppress forward locomotion (Carreira-Rosario et al., 2018). The leech R3b-1 descending neurons are command neurons involved in intersegmental coordination (Puhl et al., 2012).

The whole-body analysis of the entire connectome allowed us to identify all ascending and descending neurons and their connections in the three-day-old larva.

### Segment-specific and segmentally iterated cell types

The annotated whole-body connectome allowed us to compare the neuronal complement of the various segments of the *Platynereis* larval body. The three-day-old larva has three main trunk segments with chaeta-bearing parapodia (chaetigerous segments) and a more anterior cryptic segment (CATMAID annotation: segment_0) (Steinmetz et al., 2011). In addition, the pygidium forms the posterior-most part of the body (Starunov et al., 2015)(Figure 7A, E). The ciliary bands mark the posterior segment boundaries and *engrailed* expression the anterior boundary. Larval segments differ in the expression of Hox genes (Steinmetz et al., 2011). During the process of cephalic metamorphosis, the first segment loses its chaetae and fuses into the head (Fischer et al., 2010).

We identified many segment-specific and pygidium-specific neuron types and neuron types present in different subsets of segments (Figure 7 – figure supplement 1). There are 10 neuronal cell types specific to the first segment, eight to the second segment, two to the third segment and seven to the pygidium. Neurons with global reach are only present in the first segment, the pygidium and the head (Figure 7C-E, Figure 7 – figure supplement 1).

These distinct neuron complements suggest a functional specialisation of the different trunk segments and are in agreement with segmental differences in the expression of developmental transcription factors (Vergara et al., 2017).

There are also several neuron types that are present in all chaetigerous segments (Figure 7 – figure supplement 1) including the chaeMech sensory neurons, the INsplitPB and INbackcross interneurons and the MNche and MNhose motoneurons. In addition, three sensory neuron types and two muscle types occur in the head, in all four trunk segments and the pygidium (Figure 7F-J).

### Exocrine glands and their innervation

There are two types of non-locomotor effector systems in the *Platynereis* larva, exocrine glands and pigment cells.

Two large modules are centred around a pair of exocrine-gland motoneurons (MNspinning). These have the largest number of incoming and outgoing synapses in their respective modules (Figure 2). The two MNspinning cells synapse on two pairs of large endocrine gland cells (spinGland or spinning gland) found in the parapodia of the second and third segments.

The four spinGland cells have a large microvillar secretory pore at the tip of the ventral parapodia (neuropodia). The MNspinning motoneurons are postsynaptic to various sensory pathways, including the chaetal receptors (chaeMech), two head SNbronto sensory neurons and their postsynaptic interneuron partners (INbronto). Inputs from the chaeMech receptors suggest that mechanical stimuli to the chaetae may regulate secretion from the spinGlands.

We identified five further gland-cell types in the *Platynereis* larva. These differ in their position, size, ultrastructure and innervation (Figure 8A, Figure 8 – figure supplement 1). A similar diversity of glands occurs in the epidermis of other annelids (Hausen, 2005).

**Figure 8.**
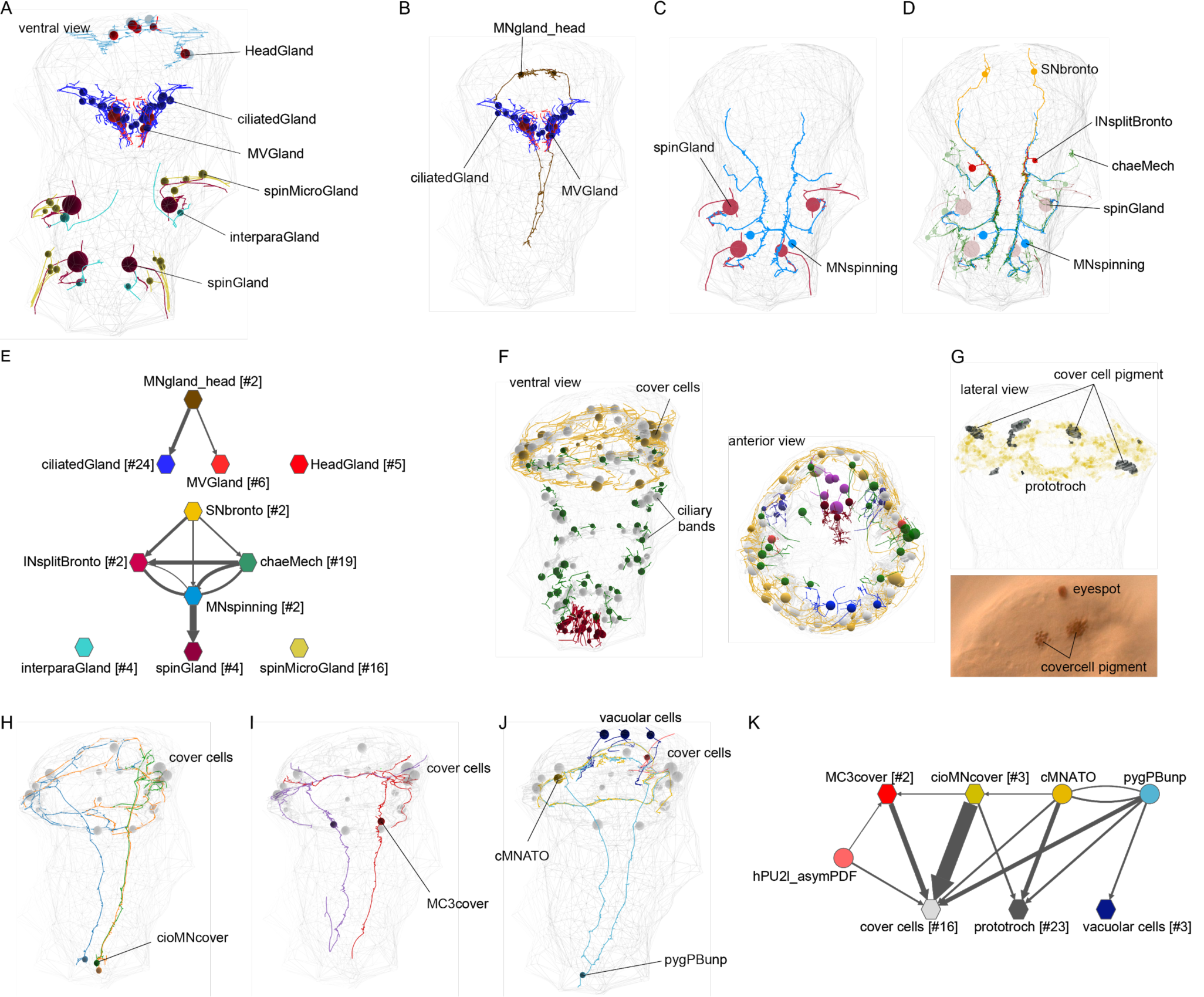
Exocrine glands and pigment cells and their innervation. (A) Overview of the position of all exocrine gland cells in the EM volume. (B) EM reconstruction of the two MNgland_head motoneurons innervating the ciliatedGland and MVgland cells. (C) EM reconstruction of the two MNspinning motoneurons innervating the four spinGland cells. (D) EM reconstruction of selected input neurons to the MNspinning cells. (E) Network view of the exocrine gland synaptic circuits. Nodes represent groups of neurons, edges represent synaptic connections. Numbers in square brackets indicate the number of cells. The headGland, interparaGlands and spinMicroGland cells receive no synaptic input. (F) Overview of the position of all pigmented cells in the EM volume, ventral and anterior views. (G) Reconstructed cover cells in the EM volume (top) and in a live larva in a DIC image (bottom) showing the position of pigment granules in the cover cell. (H) Reconstruction of the cioMNcover cells. (I) The MC3cover cells. (J) The pygPBunp cells and their postsynaptic partners, the cover cells and the head vacuolar cells. (K) Network view of the pigment cell synaptic circuits. Nodes represent groups of neurons, edges represent synaptic connections.

In the ventral head, there are five large headGland cells. These cells are filled with large (diameter=1.4 µm, stdev=0.28 N=36) secretory vesicles and have no presynaptic partners. In the first segment, there is a ventral girdle of gland cells containing two gland types, the ciliatedGland (24 cells) and the MVGland cells (6 cells). CiliatedGland cells have a microvillar collar and a stiff cilium penetrating the cuticle, suggesting that these are sensory-exocrine cells. MVGland cells have a broader microvillar secretory pore and no cilium. These gland cells are postsynaptic to two decussating head gland motoneurons (MNgland)(Figure 8B). The MNgland cells receive input from the rhythmically active serotonergic system (Verasztó et al., 2017a) suggesting a link between ciliary swimming and glandular secretion.

Close to the site of secretion of the large spinGlands, there is a secretory pore for the smaller, spinMicroGland cells (16 cells). SpinMicroGland cells have microvilli and secrete through a narrow tunnel in the cuticle. These spinMicroGlands receive no synapses in the three-day-old larva.

In the second segment, there are two additional interparapodial glands (interparaGland) with a small microvillar secretory pore opening in the cuticle between the neuro- and notopodia. These cells have long projections, but we could not identify synaptic inputs to them.

The various glands can also be distinguished by their glycosylated secretory content, as revealed by stainings with fluorescently labelled lectins (Figure 8 – figure supplement 1). The spinGland cells can be stained with wheat germ agglutinin (WGA) and *Lotus, Ulex* and *Pisum* lectins. The secretory content of headGlands can be labelled with *Griffonia*, peanut and *Pisum* lectins (Figure 8 – figure supplement 1). The girdle of ciliatedGland and MVGlandcells stains with *Griffonia* lectin (Figure 8 – figure supplement 1).

### Pigment cells and their innervation

The second type of non-locomotor effector cells comprises various pigmented epithelial cell types (Figure 8 – figure supplement 2). We distinguished seven pigmented cell types, three of them under neuronal control.

The largest and most densely innervated pigmented cells are the cover cells (16 cells). These form two rows anterior and posterior to the prototroch ciliary band. The cover cells receive synaptic innervation from three groups of neurons with global reach. The largest number of synapses are provided by three pygidial neurons (cioMNcover) with axons reaching the head. The cioMNcover axons form a double ring around the prototroch, innervating the cover cells both above and below the ciliary band (Figure 8H). The unpaired mechanosensory-pigment-motor pygPBunp neuron projects in a single ring around the cover cells and also synapse on the cover cells. The third pigment-motor cell type is the biaxonal MC3cover neuron with its soma in the first segment (2 cells). This cell has branched axons with smaller branches in between the cover cells that terminate near the cuticle. MC3cover cells are postsynaptic to the SNbronto, SNblunt and SNFVa putative mechanosensory neurons.

All other ciliary band cells are flanked by a different pigment-cell type, the CB pigment cells (58 cells). These cells contain pigment vacuoles with an average diameter of 0.34 µm (stdev=0.08, N=44). In the pygidium, there is a ring of 26 pigmented cells with larger pigment vacuoles (diameter=0.55 µm, stdev=0.14, N=47). In the head, there are three vacuolar cells with brighter vacuolar content. These cells also receive synapses from the pygPBunp neuron. The eyespots and adult eyes also have distinct types of shading pigment cells, with no synaptic partners (Jékely et al., 2008; Randel et al., 2013; Rhode, 1992).

### Chaetal mechanoreceptors and their circuits

A prominent trunk mechanosensory system with a high level of connectivity is the circuit of chaetal mechanoreceptors (chaeMech). The chaeMech neurons are dendritic sensory cells found in segments 1-3 in both the dorsal (notopodium) and ventral (neuropodium) lobes of the parapodia. Their sensory dendrites branch between the chaetal sacs and their axon projects to the nerve cord (Figure 9A-C). The sensory dendrites may sense the displacement of the chaetae during crawling (proprioception) or due to external mechanical stimuli. The annelids *Harmothoë* (a polynoid) and *Nereis* have cells with a similar morphology called bristle receptors. These cells show rapidly adapting spikes upon the displacement of the chaetae (Horridge, 1963); (Dorsett, 1964). The chaeMech cells are also similar to dendritic proprioceptors in *Drosophila* larvae that sense body-wall deformations and provide feedback about body position through premotor neurons (He et al., 2019; Vaadia et al., 2019; Zarin et al., 2019).

**Figure 9.**
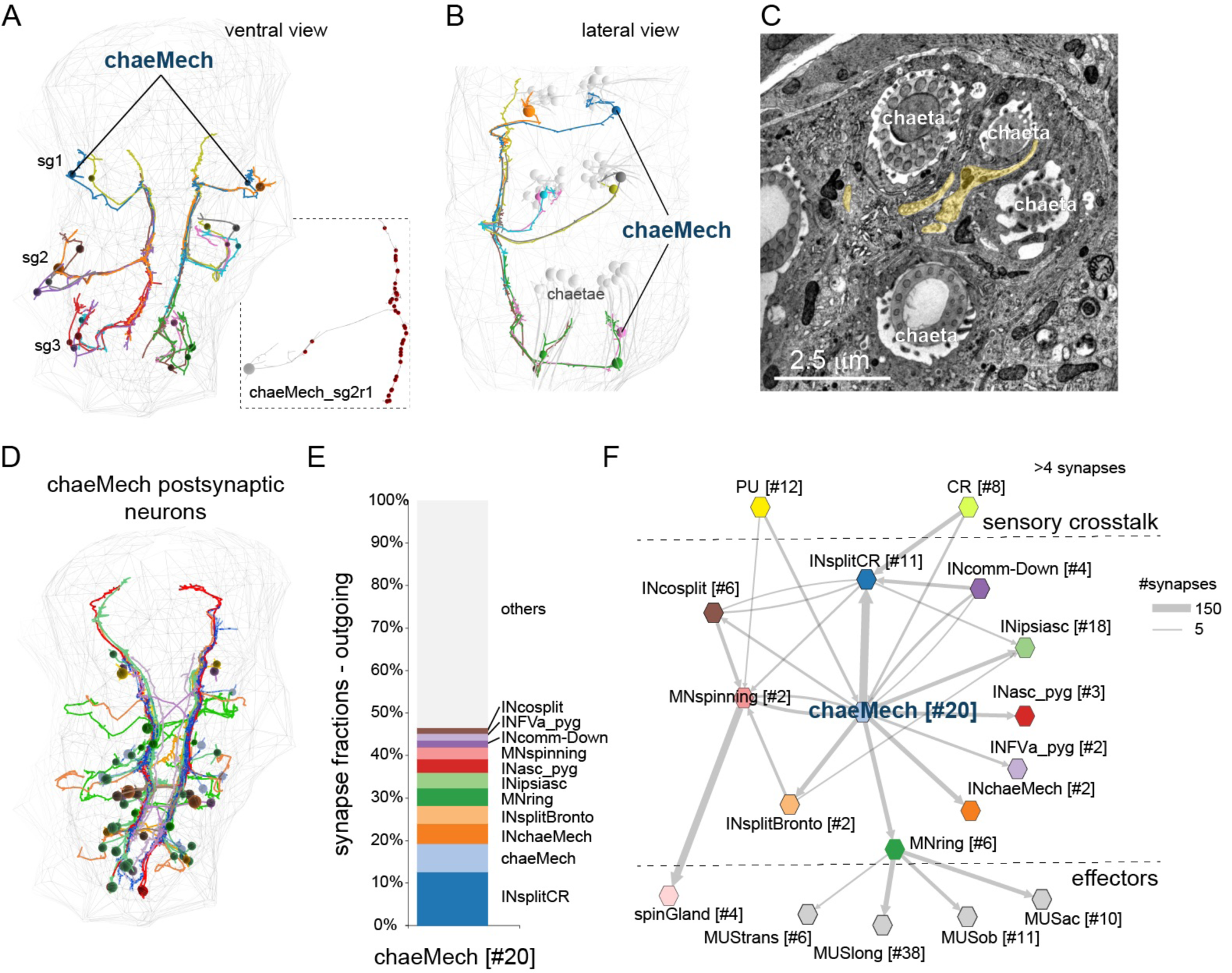
Chaetal mechanoreceptors and their circuits. (A) Skeletons of chaeMech cells in segments 1-3. Inset shows the pseudounipolar morphology of one of the cells. Red dots indicate presynaptic sites. (B) Lateral view of chaeMech cells. The branched dendritic endings run in between the chaetae in the chaetal sacs. (C) TEM image of a chaetal sac with cross-sections of chaetae and chaeMech dendrites (highlighted in yellow). (D) Skeletons of the main postsynaptic targets of chaeMech cells. (E) Fraction of outgoing synapses of chaeMech cells to their main postsynaptic partners. (F) Grouped connectivity graph of chaeMech cells. Nodes represent groups of cells, arrows represent synaptic connections. Edge width shown as the square root of synaptic count.

In *Harmothoë* and *Nereis*, the bristle receptors connect to the giant axon system. The *Platynereis* chaeMech neurons are highly interconnected. They form some of the strongest edges in the connectome (Supplementary table 1) and have the highest number of direct postsynaptic targets (Figure 9D-E; Figure 5A, C). These include the premotor interneurons INsplitCR, INsplitBronto, INchaeMech and the MNring and MNspinning motoneurons. Some trunk PU and CR mechanoreceptors (Bezares-Calderón et al., 2018) are presynaptic to chaeMechs, suggesting a crosstalk between different mechanosensory modalities. The INsplitCR interneurons receive the largest fraction of chaeMech synapses (Figure 5E). INsplitCRs are also postsynaptic to the collar receptors (CR) that mediate a hydrodynamic startle response characterised by parapodial extension and chaetal opening (Bezares-Calderón et al., 2018). This wiring suggests that during a startle, proprioceptive feedback into INsplits by chaeMech cells may regulate the response.

### Mechanosensory circuits and circuit evolution by duplication

The various mechanosensory systems across the body show similarities in their cellular complements and anatomy (Figure 10). There are several mechanosensory cell types in *Platynereis* larvae, as supported by various lines of evidence. Collar receptor (CR) neurons have a penetrating sensory cilium surrounded by a collar of 10 microvilli. CR neurons express the polycystins *PKD1-1* and *PKD2-1*, and detect water-borne vibrations. PB (penetrating biciliated) neurons have two penetrating cilia with a collar and express *PKD2-1* but not *PKD1-1* (Bezares-Calderón et al., 2018)(Figure 10A, B). PU (penetrating uniciliated) neurons have one penetrating sensory cilium surrounded by a collar. These neurons express the mechanosensory channel *NompC* (LABC and GJ, unpublished). InterparaPM (penetrating multiciliary) neurons have 3-4 penetrating cilia, suggesting that they are also mechanosensory (Figure 10A, B). The chaeMech cells are dendritic chaetal mechanoreceptors (Figure 9A-C).

**Figure 10.**
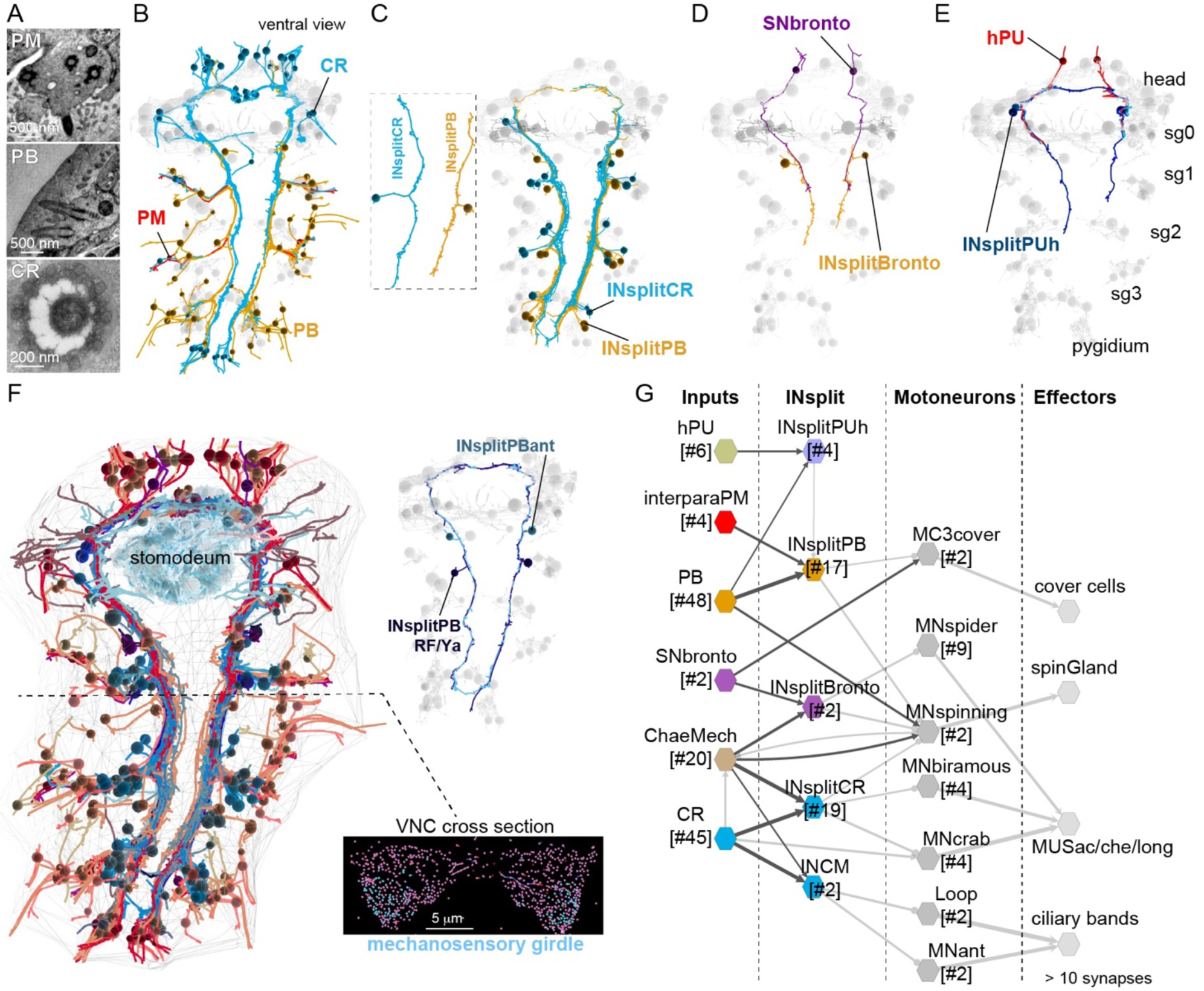
Cells and circuits of the mechanosensory girdle. (A) TEM images of the sensory endings of interparaPM, PB and CR neurons. (B) Skeletons of CR, PB and interparaPM (PM) neurons. (C) Skeletons of INsplitCR and INsplitPB neurons. (D) Skeletons of SNbronto and INsplitBronto neurons. (E) Skeletons of INsplitPUh and two hPU neurons. (F) Skeletons of all neurons projecting into the mechanosensory girdle. Upper inset: the INsplitPBant and INsplitPB_RF/Ya neurons outline the mechanosensory girdle. Lower inset: cross-section of the ventral nerve cord (VNC) indicating the position of neurites of the girdle (cyan). (G) Synaptic connectivity graph of the main sensory and interneuron types of the mechanosensory girdle. Their main motoneuron targets and effector systems are also shown. Edges represent synaptic connections, nodes represent groups of cells.

The strongest postsynaptic partners for all these mechanoreceptors are ipsilateral interneurons with a similar bifurcating axon. These interneurons – collectively referred to as INsplit – have several distinct types (INsplitCR, INsplitPB, INsplitPBant, INsplitPB_RF/Ya, INsplitPUh, INsplitBronto, INCM)(Figure 10C-E). INsplit neurons occur in all four trunk segments and in the head (INsplitPUh) and their distinct types have unique synaptic connectivity. PB and interparaPM neurons specifically target INsplitPB and represent their main input. The dendritic chaeMech and SNbronto neurons both synapse on the INsplitBronto interneurons, which have no other major presynaptic partners.

CR neurons synapse on INsplitCR and INCM. ChaeMech cells also strongly target INsplitCR. Some head PU neurons synapse on INsplitPUh.

The repeated pairing of distinct mechanoreceptor cell types with distinct INsplit types suggests an evolutionary pathway for these systems by circuit duplication and divergence.

### The mechanosensory girdle

The mechanosensory circuits in the larva collectively outline an anatomical system that we call the mechanosensory girdle. The girdle is characterised by a circular axonal track with a dorsal loop in the brain neuropil and two axon bundles along the trunk ventral nerve cord (VNC), connected at the circumesophageal nerve (Figure 10F, Video 3). The global projections of two INsplit types – INsplitPBant and INsplitPB_RF/Ya – outline the axon tracks of the mechanosensory girdle (Figure 10F). The projections of several other neurons follow this axonal track and form a bundle at the ventral side of the VNC (Figure 10F). The projections of the CR, PB, PU, interparaPM, chaeMech sensory neurons and of all INsplit interneurons run along the mechanosensory girdle, suggesting a separate VNC track for the processing of mechanosensory signals. Searching for pre- and postsynaptic partners for all these neurons we identified further neurons that are part of this system. These include four further sensory cell types, the head SNbronto (dendritic) and SNblunt neurons (blunt sensory ending beneath the cuticle, no cilium) and two pygidial cell types, SNPDF_pyg and SNpygM (Figure 10 – figure supplement 1). Additional interneuron types that join the girdle are the pygidial ascending INasc_pyg (Figure 4D) and the head descending INrope neurons (Bezares-Calderón et al., 2018). The ciliomotor neurons MNant (Verasztó et al., 2017a) and the cover cell motoneurons MC3cover (Figure 5E) are also part of the mechanosensory girdle.

This anatomy and the shared circuitry suggest that the various mechanosensory modalities are integrated by the mechanosensory girdle system. Overall, mechanosensory neurons provide input to all effector systems, including muscles, ciliated cells, glands and pigment cells (Figure 10G).

## Discussion

### The *Platynereis* larval connectome and cell-type atlas

The *Platynereis* three-day-old larva connectome is the third and currently largest whole-body connectome, after those of *Caenorhabditis elegans* and the *Ciona intestinalis* larva (Cook et al., 2019; Ryan et al., 2016; White et al., 1986). To facilitate the browsing and querying of the *Platynereis* connectome, the entire dataset – including EM images, skeletons, tags, connectors and annotations – will be available at https://catmaid.jekelylab.ex.ac.uk. We also provide R scripts for data retrieval from the CATMAID server by libraries in the `natverse` toolkit.

The connectome will allow the generation of specific circuit-level hypotheses about neuronal control in the *Platynereis* larva. For example, it suggests that mechanosensory circuits regulate exocrine glands and pigments cells. We also identified several potential cases of multisensory integration. For example, the eyespot R1 and adult eye photoreceptors converge on the INR1 interneurons suggesting the integration of directional light sensing between the ventral eyespots and dorsal adult eyes.

The EM volume also allowed us to classify all neuronal and non-neuronal cells into 270 types. It will be interesting to compare this morphology and connectivity-based classification to single-cell RNAseq-based classifications. We expect a similar number of molecular types, as suggested by the already available molecular data for *Platynereis (Achim et al*., *2015)*, and also by the similar number of morphological and molecular cell types in both the *Ciona* larva and *Caenorhabditis elegans* (Cao et al., 2019; Hobert et al., 2016).

### Limitations of the connectome

There are several limitations of connectomics. In *Platynereis*, for example, we know the neurotransmitter or neuromodulator content only of a small fraction of neurons (Figure 4) as these molecules are invisible to EM. However, artificial convolutional neural networks were recently used to predict synaptic transmitter content in the *Drosophila* adult brain from EM images alone (Eckstein et al., 2020). This approach still has to be tested in other organisms including *Platynereis*. An alternative is to register cellular-resolution gene-expression atlases to EM volumes, as has recently been done for six-day-old *Platynereis* (Vergara et al., 2020). The mapping of neurotransmitter-synthesis enzymes or transporters could then reveal transmitter content in single neurons. This is now feasible in *Platynereis* larvae but will require the acquisition of new synaptic-resolution EM volumes. We attempted but could not register the current volume to a three-day-old gene expression map (Asadulina et al., 2012) due to differences in fixation and alignment artefacts.

Another approach to map transmitters and modulators in small and stereotypical nervous systems is to correlate transgenically labelled neurons to EM reconstructions, an approach extensively used in *C. elegans* (Bentley et al., 2016) and also possible in *Platynereis* larvae (Bezares-Calderón et al., 2018; Verasztó et al., 2017a). Alternatively, we have used direct immunogold labelling on sparse sections from the EM series to directly map neuropeptides to the connectome (Shahidi et al., 2015)(Figure 4).

Another limitation of the *Platynereis* connectome is the difficulty to predict sensory-cell function based on ultrastructure alone. We identified 45 sensory neuron types in the larva based on morphology and connectivity, but only know the sensory function of a few of these (Williams and Jékely, 2019). However, it is now possible to combine genetics, behaviour and neuronal activity imaging in *Platynereis* to functionally characterise sensory cell types. Recent work has identified larval UV photoreceptors (Verasztó et al., 2018), hydrodynamic vibration detectors (Bezares-Calderón et al., 2018), chemosensory neurons (Chartier et al., 2018), or the Go-opsin1-expressing shadow detectors in sensory appendages of adult worms (Ayers et al., 2018). The connectome nevertheless allowed us to identify new types of putative mechanosensory neurons – including the SNblunt, SNFVa and SNbronto cells – based on connectivity and morphology alone (Figure 9).

### Circuits for whole-body coordination

*Platynereis* larvae have segmentally repeated effector organs including ciliary bands, parapodia and glands. The connectome suggests that for each of these effector systems, neurons with axons spanning several segments can ensure intersegmental coordination.

Several motoneurons innervate target cells in multiple segments. The two MNspinning neurons innervate the four spinning glands in segments 2-3 with each MNspinning connecting to two glands on the contralateral body side (Figure 8C). Among the muscle motoneurons, MNspider cells synapse on contralateral muscles in two consecutive segments. The muscle targets of MNcrab and MNbow neurons span three segments (Bezares-Calderón et al., 2018). In the pigment-motor system, the cioMNcover cells each innervate half the ring of the cover cells and pygPBunp innervates the entire ring, suggesting coordinated regulation. The ciliomotor system is characterised by large motoneurons coordinating entire ciliary bands across all segments (Verasztó et al., 2017a). These intersegmental motor systems likely mediate behaviours including the simultaneous recruitment of multiple effectors. Those behaviours that involve intersegmental phase lags such as parapodial crawling (not fully developed at three days) may be mediated by segment-intrinsic motoneurons (e.g. MNantacic, MNpostacic, MNpostv).

Intersegmental coordination is also apparent at the level of sensory and interneurons. For example, individual chaeMech cells can synapse on interneurons in three segments. INrope and INchaeMech interneurons target motoneurons across three segments, INsplitCRATO, INsplitBronto and INsplitVent in two segments. The globally reaching INsplitPBant cells have postsynaptic partners in all four segments and the pygidium.

These examples demonstrate the importance of a whole-body approach in connectomics for the identification of such long-range connections.

### Circuit evolution by duplication and divergence

The connectome suggests a model for the evolution of circuits by the duplication and divergence of circuit modules of synaptically connected cell types (Tosches, 2017). This was most apparent in the mechanosensory system where several morphologically similar collared ciliated mechanosensory neurons (CR, PU, PB, PM) synapse on distinct groups of morphologically similar INsplit types.

The similar morphologies of the mechanosensory cells suggests that these represent sister cell types (Arendt, 2008). Further supporting this, PB, CR and pygPBunp neurons share some of their gene regulatory environment as they all can be labelled by a *PKD2-1* transgene (Bezares-Calderón et al., 2018). The cell-type diversification of mechanoreceptors may have been paralleled by the diversification of the postsynaptic INsplit neurons. The duplication and divergence of entire cell type sets has also been proposed for the evolution of the vertebrate cerebellum (Kebschull et al., 2020). Testing this model in *Platynereis* would require the integration of connectomics and comprehensive gene expression analysis.

### Segmental organisation and inferences about the evolution of the annelid body plan

The whole-body connectome allowed us to investigate the segmental organisation of the annelid body. Each larval segment has a distinct neuron-type composition with some unique cell types (Figure 7 – figure supplement 1). Several neuron types occur in multiple segments and those shared between all four trunk segments confirm the serial homology of the cryptic segment with the other three segments (Steinmetz et al., 2011). The pygidium has also been hypothesized to have had a metameric origin based on the presence of a pygidial coelomic cavity and muscles (Starunov et al., 2015). One prediction of this model is that there are neurons or muscles of the same type in the pygidium and the main trunk segments. We identified four neuronal (trunk PB, INipsiasc, trunk CR and PU) and two muscle types (MNlong_V, MNtrans) showing this pattern (Figure 7), supporting the metamery hypothesis.

The identification of a mechanosensory girdle in *Platynereis* reminded us of another classic hypothesis about bilaterian body plan evolution, the amphistomy theory. In 1884, Sedgwick suggested that at the origin of bilaterians, a gastric slit in a radially symmetric animal evolved into the mouth and anus through the fusion of the lateral lips (Sedgwick, 1884). If there was a nerve concentration around the gastric opening, like in some cnidarians, this could have evolved into the paired nerve chords characteristic of many bilaterians (Nielsen et al., 2018). Accordingly, the mechanosensory cells in the *Platynereis* girdle could correspond to mechanosensory cells around the cnidarian oral opening (e.g. (Singla, 1975)). One prediction of this scenario is the presence of bifurcating mechanosensory interneurons similar to INsplits running in two directions along the oral opening in cnidarians. This specific neuroanatomical prediction could be tested by connectomics in cnidarians.

## Methods

### Specimen preparation, transmission electron microscopy and image processing

Fixation and embedding were carried out on an 72 hpf *Platynereis* larva (HT9-4) as described previously (Conzelmann et al., 2013). Serial sectioning and transmission electron microscopy were done as described in (Shahidi et al., 2015). The section statistics for the HT9-4 (NAOMI) specimen were previously described (Randel et al., 2015).

Serial sections were imaged on a FEI TECNAI Spirit transmission electron microscope with an UltraScan 4000 4×4k digital camera using Digital Micrograph acquisition software (Gatan Software Team Inc., Pleasanton) and SerialEM (Schorb et al., 2019). The images for the HT9-4 projects were scanned at various pixel resolutions: 5.7 nm/pixel, 3.7 nm/pixel, and 2.2 nm/pixel. Image stitching and alignment were done in TrakEM2 (Cardona et al., 2012). We used CATMAID for skeleton tracing, reviewing, annotation and connectivity analysis (Saalfeld et al., 2009; Schneider-Mizell et al., 2016).

Due to contrast and focus problems in the main dataset we had to re-image certain layers at higher resolutions, to allow tracing of neurons. These re-imaged series were made into independent CATMAID projects. This included five extra projects, taken at various points throughout the main dataset (Table 4). The largest of these, Plexus_HT-4_Naomi_project__372-4013, consisted of 1407 layers at resolution of 2.2 nm. This stack mostly focused on the brain plexus and the ventral nerve cord where most neurites and synapses occur. Other projects consisted of three jump/gap regions that required not only high resolution but also better realignment. One set contained all the immunogold labelled layers (Shahidi et al., 2015) that were not included in the main aligned dataset. These layers had very low contrast due to the immunolabelling procedure and therefore required higher resolution imaging. All projects were first created and processed in TrakEM2 and then exported as flat jpeg images into CATMAID.

### Neuron tracing, synapse annotation and reviewing

To digitally reconstruct every neuron in the serial TEM dataset of the three-day-old larva, we used the collaborative web application CATMAID (Schneider-Mizell et al., 2016) installed on a local server. Ultrastructural features (number and orientation of microtubules, electron density of the cytoplasm and vesicles, ER structure) and a high resolution dataset of the neuropil and the ventral nerve cord aided tracing. At the approximate centre of each nucleus, we changed the radius of a single node according to the soma size in that layer. Skeletons were rooted on the soma and the node was tagged with ‘soma’. We identified synapses based on a vesicle cloud close to the plasma membrane. Most synapses were visible in consecutive layers (for example images see (Randel et al., 2014) and browse the data). We also checked for the proximity of mitochondria in the same arbor in case of an ambiguous synapse – a requirement supported by quantitative connectomic data in *Drosophila* (Schneider-Mizell et al., 2016).

The systematic review of all neurons belonging to a cell type was done by one or multiple reviewers until close to 100% was reached for every cell. Cells were further checked in the 3D widget to split implausible skeletons. Synapses were reviewed multiple times, from both the pre- and postsynaptic arbor.

### Cell nomenclature and annotations

All cells have a unique name. We named neurons based on their type (e.g. sensory or motor) cell body position (left or right, head or trunk segment), axonal morphology, neuropeptide expression, and other specialisations (e.g. sensory morphology). Cells of the same type have similar names, distinguished by body position indicators and numbers (Tables 1 and 2). We endeavoured to give names that were easy to remember. The name of many sensory neurons start with SN followed by a specific term (e.g. blunt, bronto, stiff). Interneurons often start with IN and motoneurons with MN. There are exceptions, including neurons with known function (e.g. PRC for photoreceptor cells) and neurons with prominent morphology (e.g. Loop). Segmental position (sg0-3) and body side (l or r) is indicated in the name of most neurons. Non-neuronal cells were named based on anatomical terms (e.g. prototroch) or by abbreviations (e.g. EC for epithelial cell).

All cells have multiple annotations, which can be used to query the database in CATMAID or in the `natverse`. Neurons belonging to a cell type category were annotated with the generalist annotation ‘celltype’ and a cell-type-specific annotation (e.g. celltype23 for the INpreMN neurons). Non-neuronal cells belonging to a cell type category were annotated with the generalist annotation ‘celltype_non_neuronal’ and a cell-type-specific annotation (e.g. celltype_non_neuronal23 for the acicula cells). Neurons were also annotated with descriptors of their projection morphologies (e.g. commissural, ipsilateral, pseudounipolar etc.), neuron class (Sensory neuron, sensory-motor neuron, sensory-neurosecretory neuron, interneuron, inter-motorneuron and motorneuron [note the ‘r’]) as defined earlier (Williams and Jékely, 2019). In CATMAID, we recommend the use of regular expressions for searching annotations e.g. ^motorneuron$ to retrieve exact matches. Differentiating neurons with immature sensory dendrites or axonal projections with axonal growth cones and with no or few synapses were annotated ‘immature neuron’ (389 cells). Ascending trunk neurons and descending head neurons traversing the circumesophageal connectives were annotated ‘head-trunk’. Neurons with a soma in the head and a descending decussating axon were annotated with ‘decussating’. Cells were also annotated according to the location of their soma in a certain body region (head – as ‘episphere’, trunk – as ‘torso’, ‘pygidium’), body side (‘left_side’, ‘right_side’), segment (‘segment_0’ etc.), and germ layer (ecto-, meso-, endoderm).

### Criteria for including cells in the connectome

We defined the final set of cells that were included in the connectome based on the number of incoming and outgoing synapses per cell. First, we selected all cells with a soma (8,852). Next, we removed all cells without a synaptic partner with a soma, resulting in 3,332 cells. We further removed cells with <3 outgoing synapses and no incoming synapse (weighted in-degree=0, weighted out-degree<3), except sensory neurons and neurons that belonged to a cell type with other well-connected members. We also removed cells with no outgoing synapses and <3 incoming synapses (weighted in-degree<3, weighted out-degree=0), except effector cells and neurons that belonged to a cell type with other well-connected members. The removed cells are ‘dead ends’ and unless they are sensory cells with output or effector cells with input, will not contribute to global connectivity. We also removed small clusters of only locally connected neurons. This resulted in a set of 2,728 cells (CATMAID annotation ‘connectome’; Figure1 – source data 1).

### Network analysis, NBLAST and Sholl analysis

We used CATMAID (several releases), Gephi 0.9.2 and Python for network analysis. To detect communities, we used Gephi with the ‘modularity’ function, considering edge weights and randomisation.

NBLAST was carried out in the `natverse` (Bates et al., 2020; Costa et al., 2016) by a custom R script. We compared 1,772 neurons in the connectome (annotated: “connectome_neuron”). First, skeletons were imported from the CATMAID database to R with the ‘read.neurons.catmaid’ command. Skeletons were smoothed by ‘smooth_neuron’ with a sigma of 6000. Skeletons were converted to point vector format by ‘dotprops’. We calculated all by all NBLAST scores by ‘nblast_allbyall’ (version 1) and clustered the results with ‘nhclust’ with the ‘ward.D2’ method.

Sholl analysis was done in CATMAID in the ‘Morphology plot’ widget (Schneider-Mizell et al., 2016). We used a radius of 100 nm and centred the analysis on the root node (soma). The Scholl profiles for the 772 connectome neurons were exported as a .csv file and analysed in R. Pairwise distances were calculated with the ‘minkowski’ method and the distance matrix was clustered with the ‘ward.D2’ method. The dendrogram was plotted with the ‘dendextend’ library. The dendrogram based on connectivity was calculated from the synaptic matrix of the 1,772 connectome neurons as presynaptic partners (rows) and the connectome cells as postsynaptic partners (columns).

## Tables

### Neuronal cell types

**Table 1.**
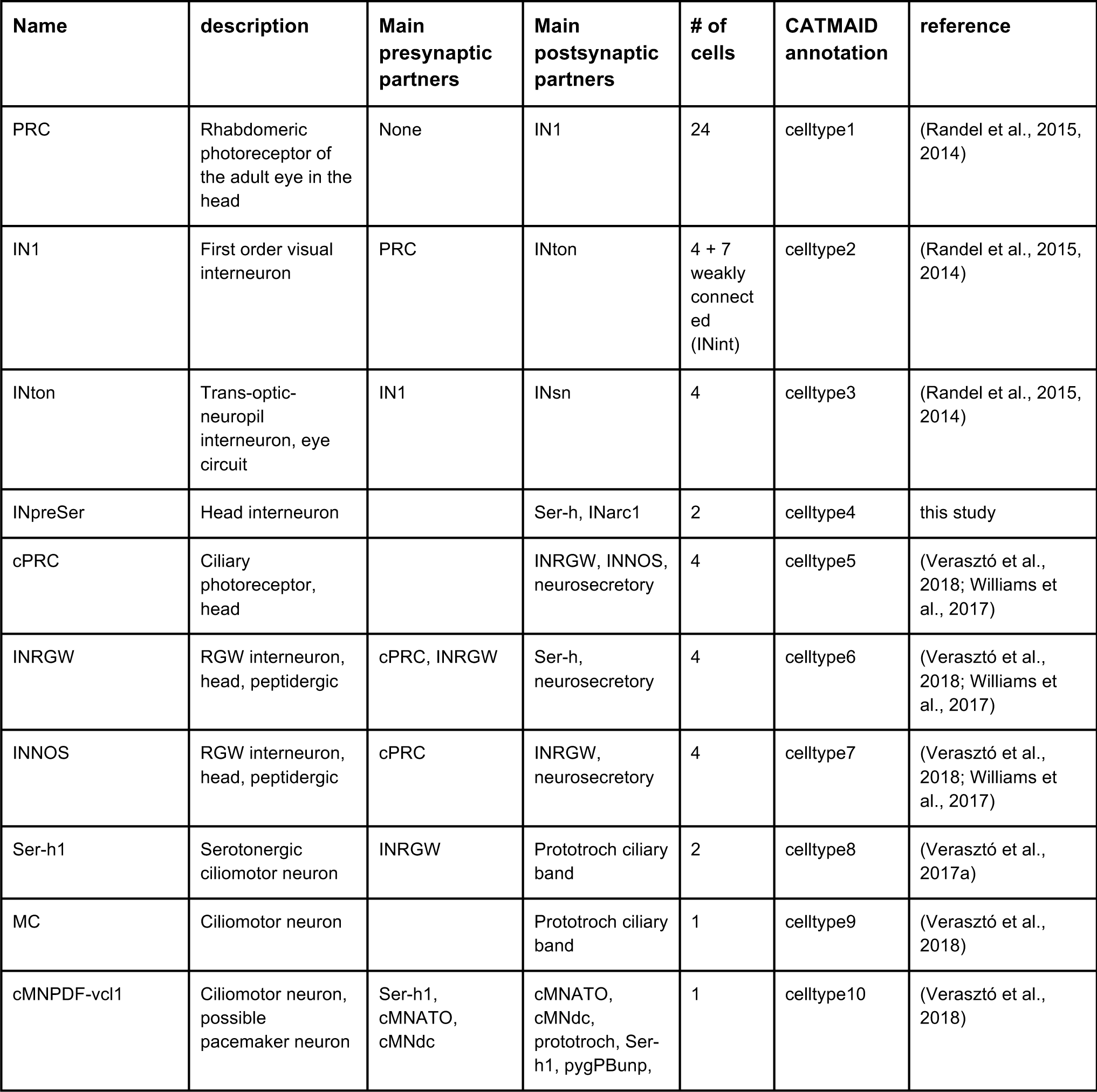
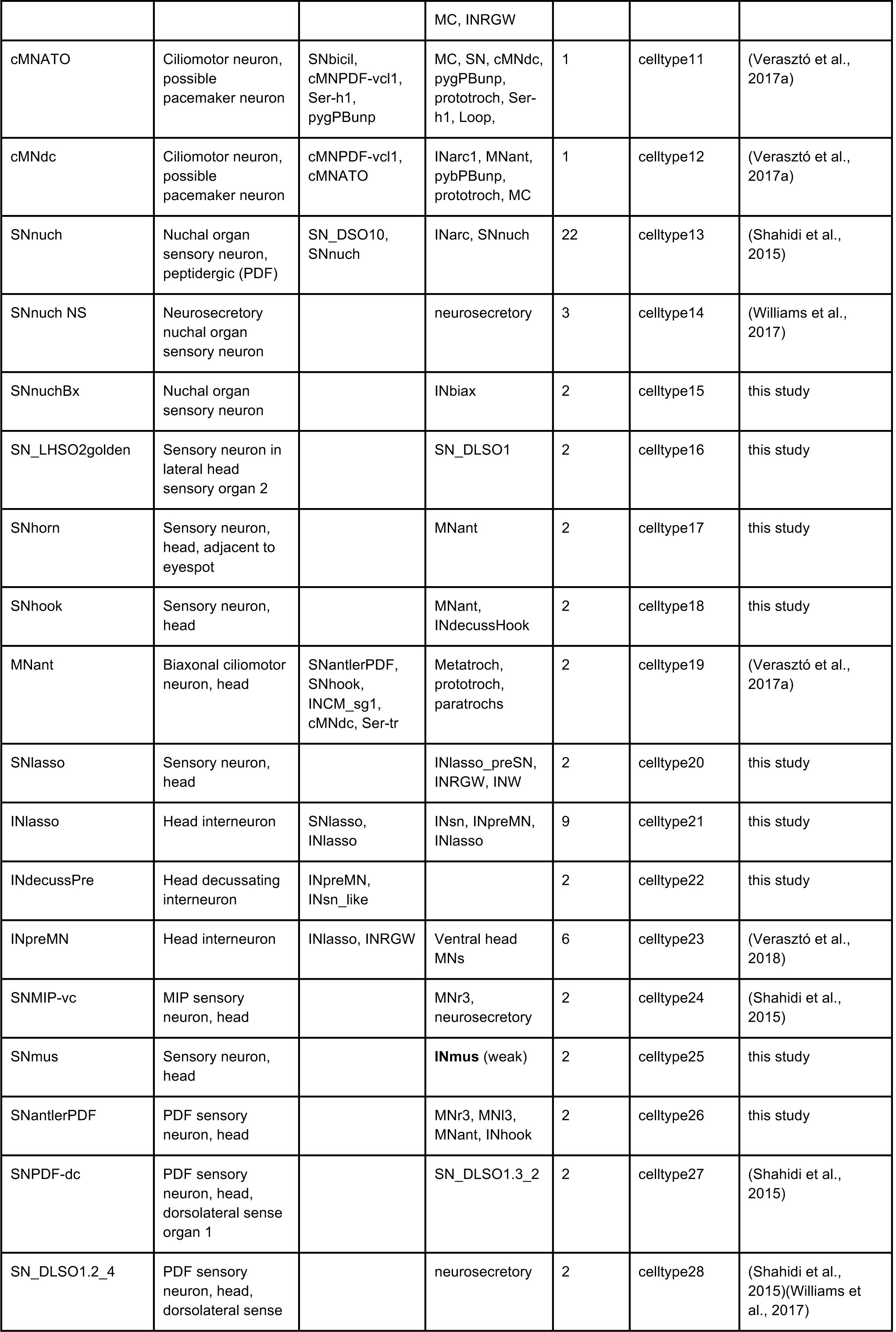
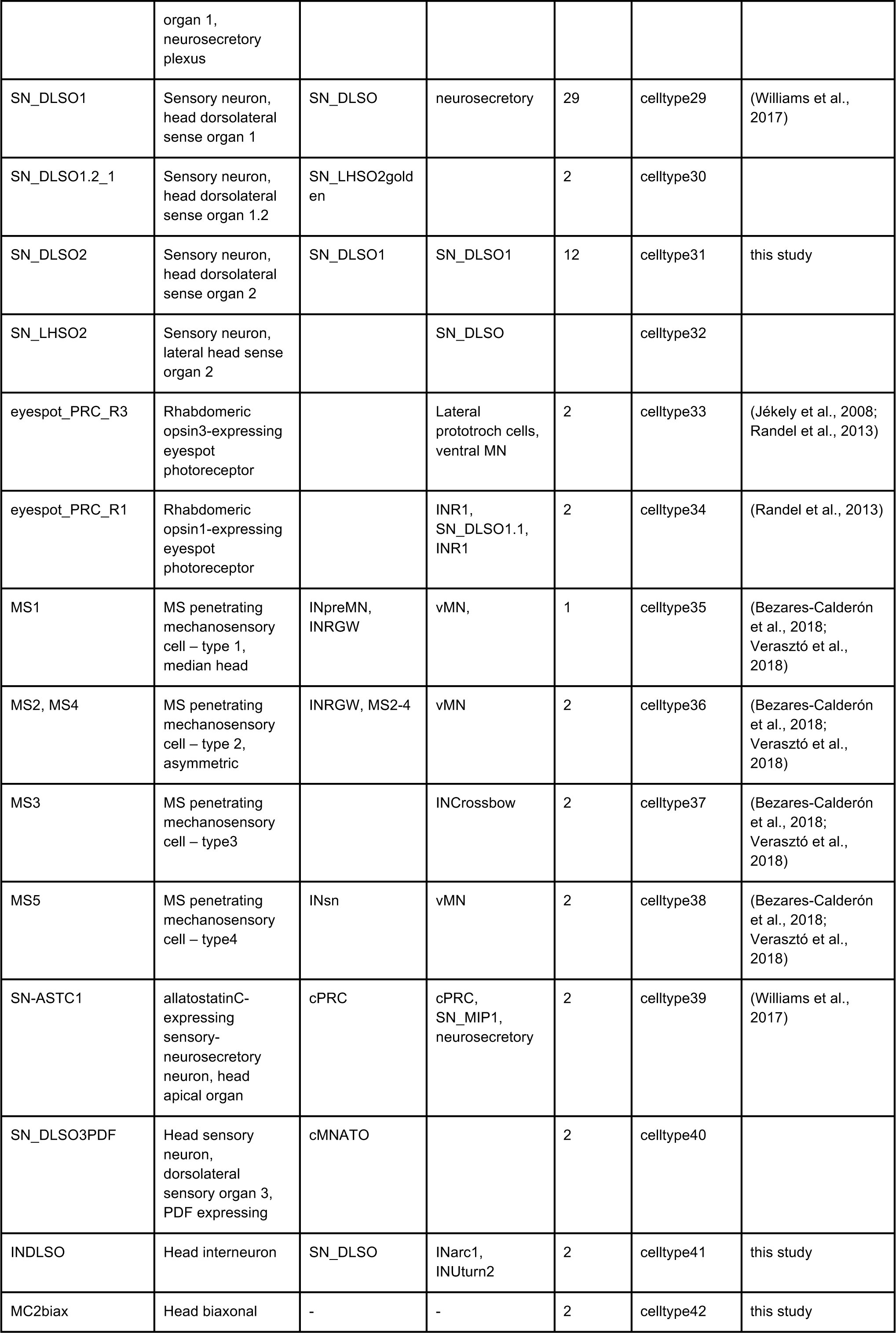
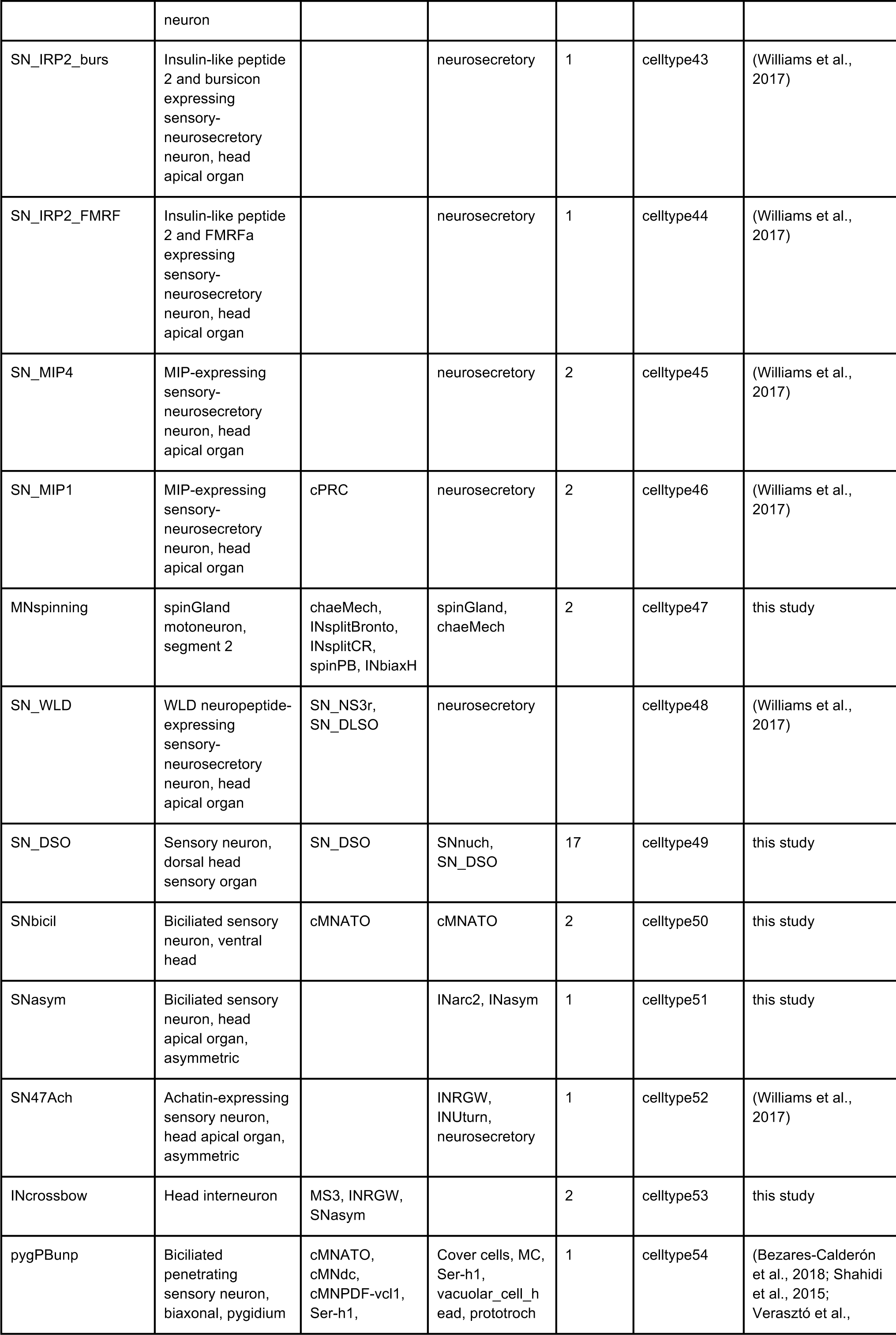
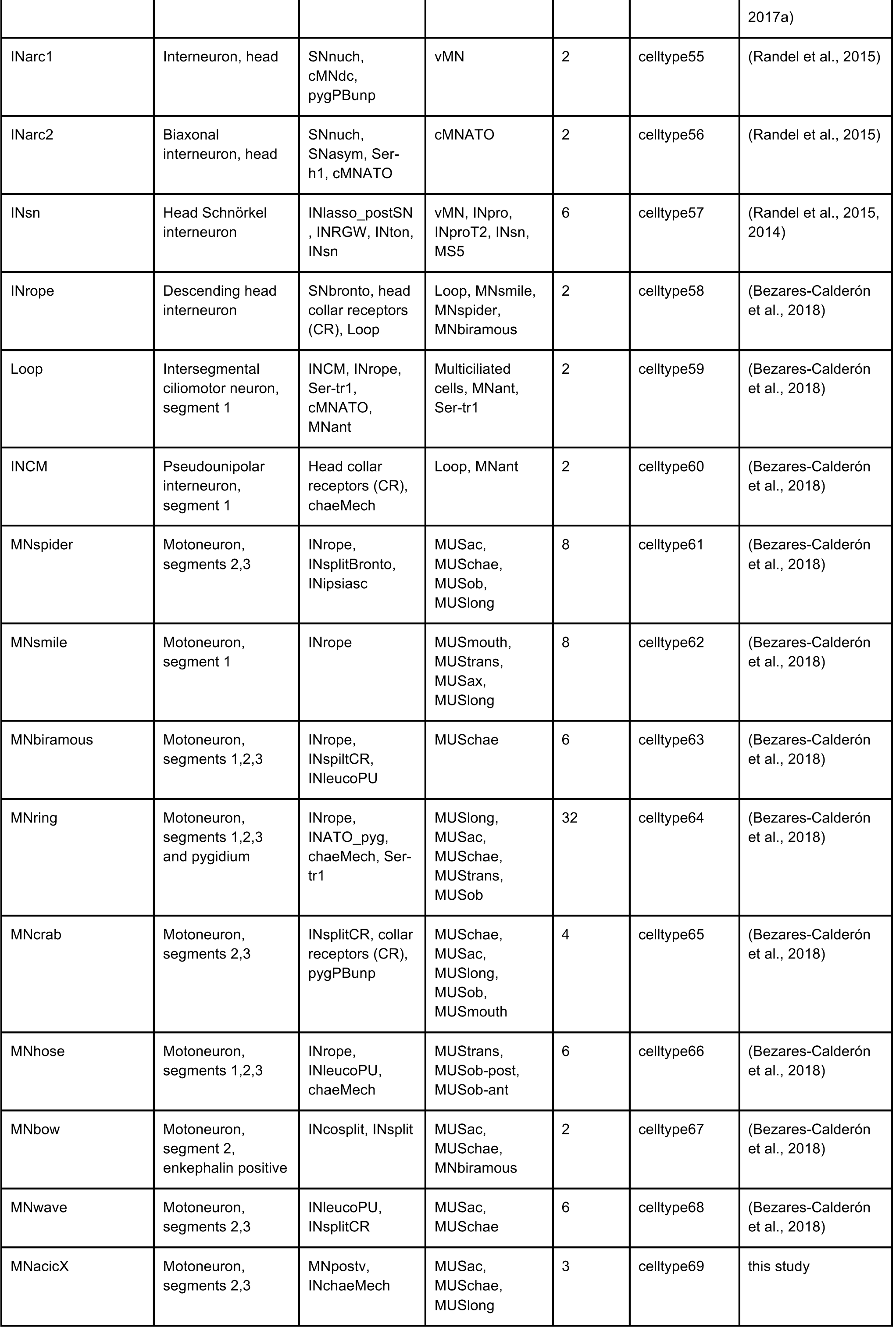
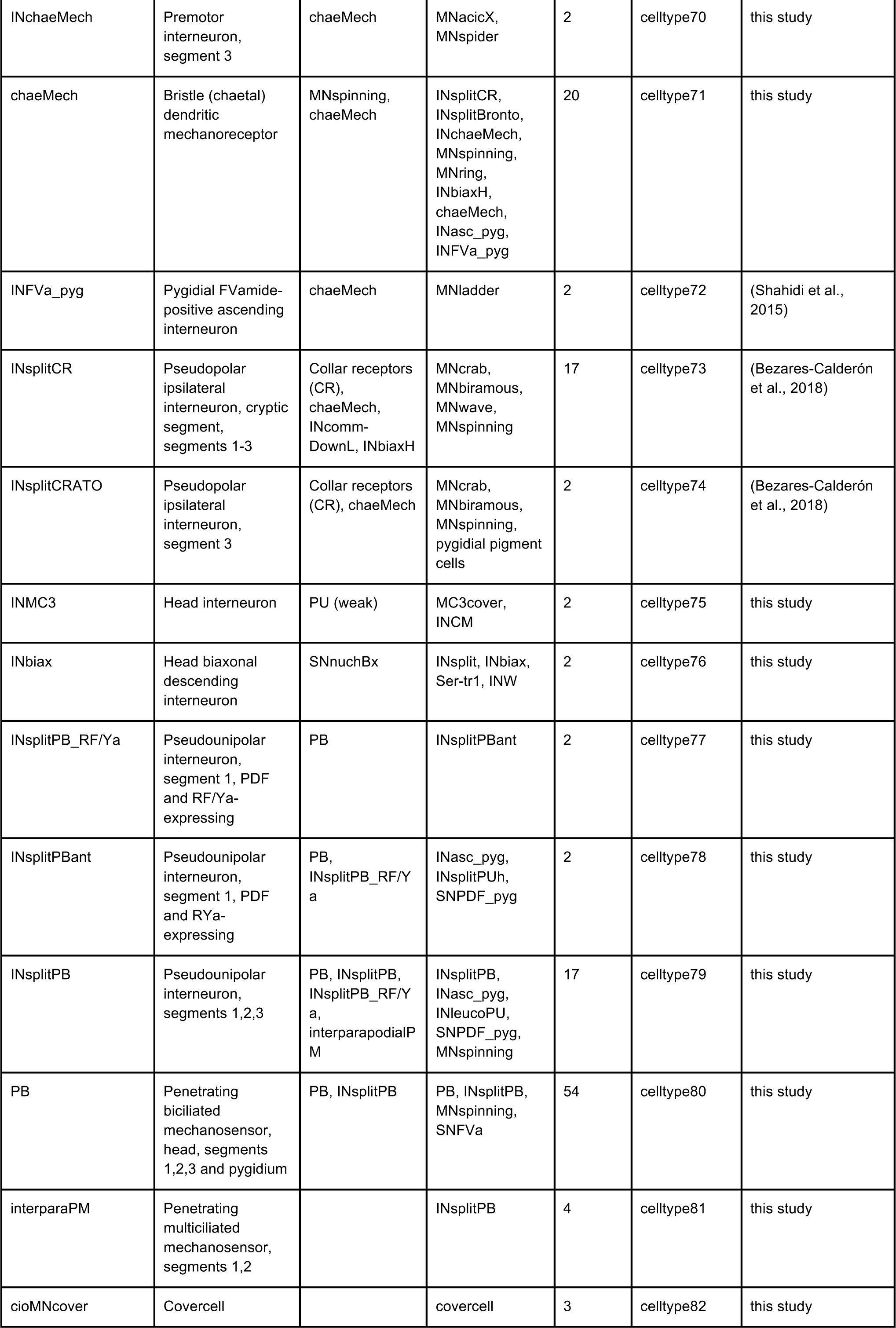
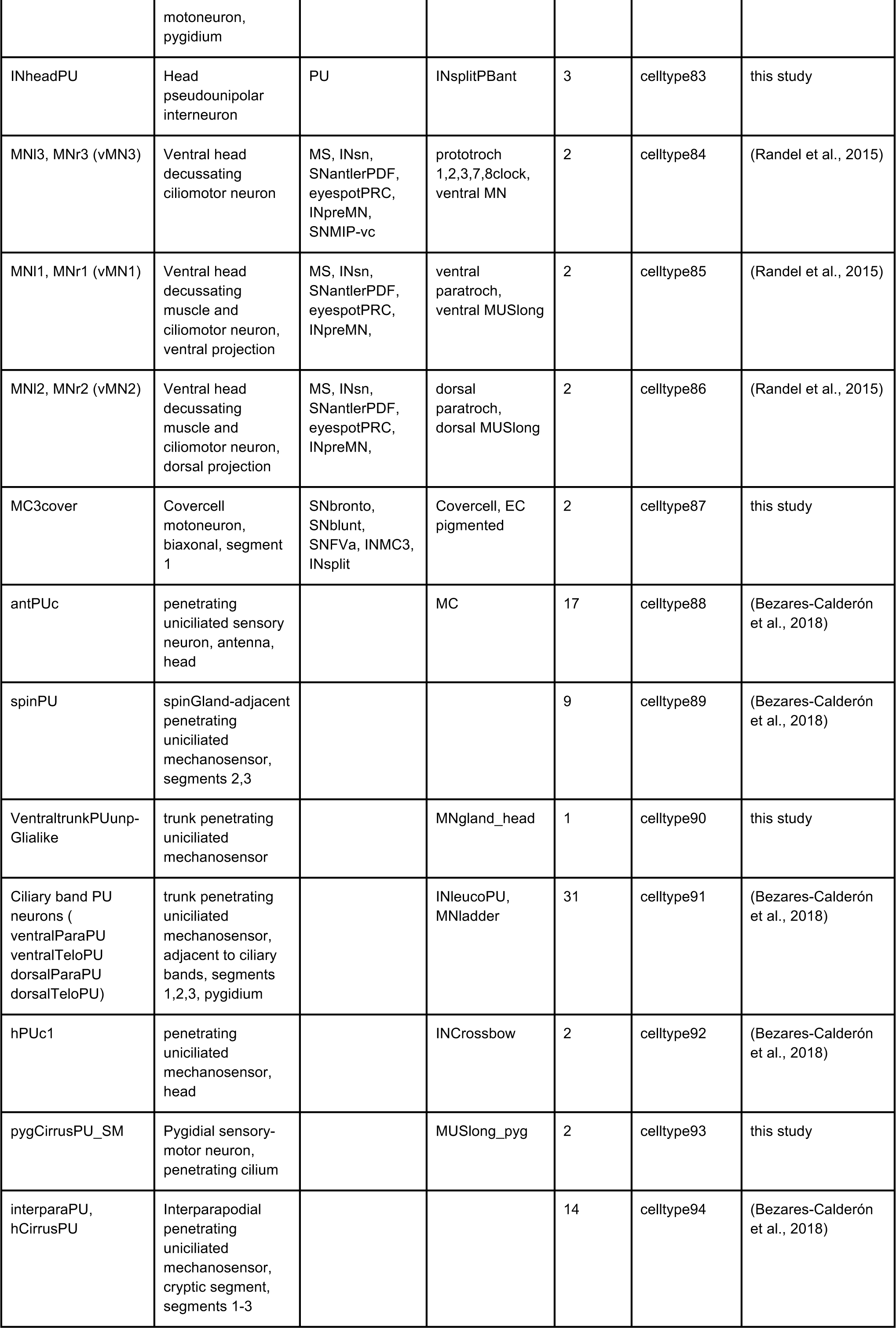
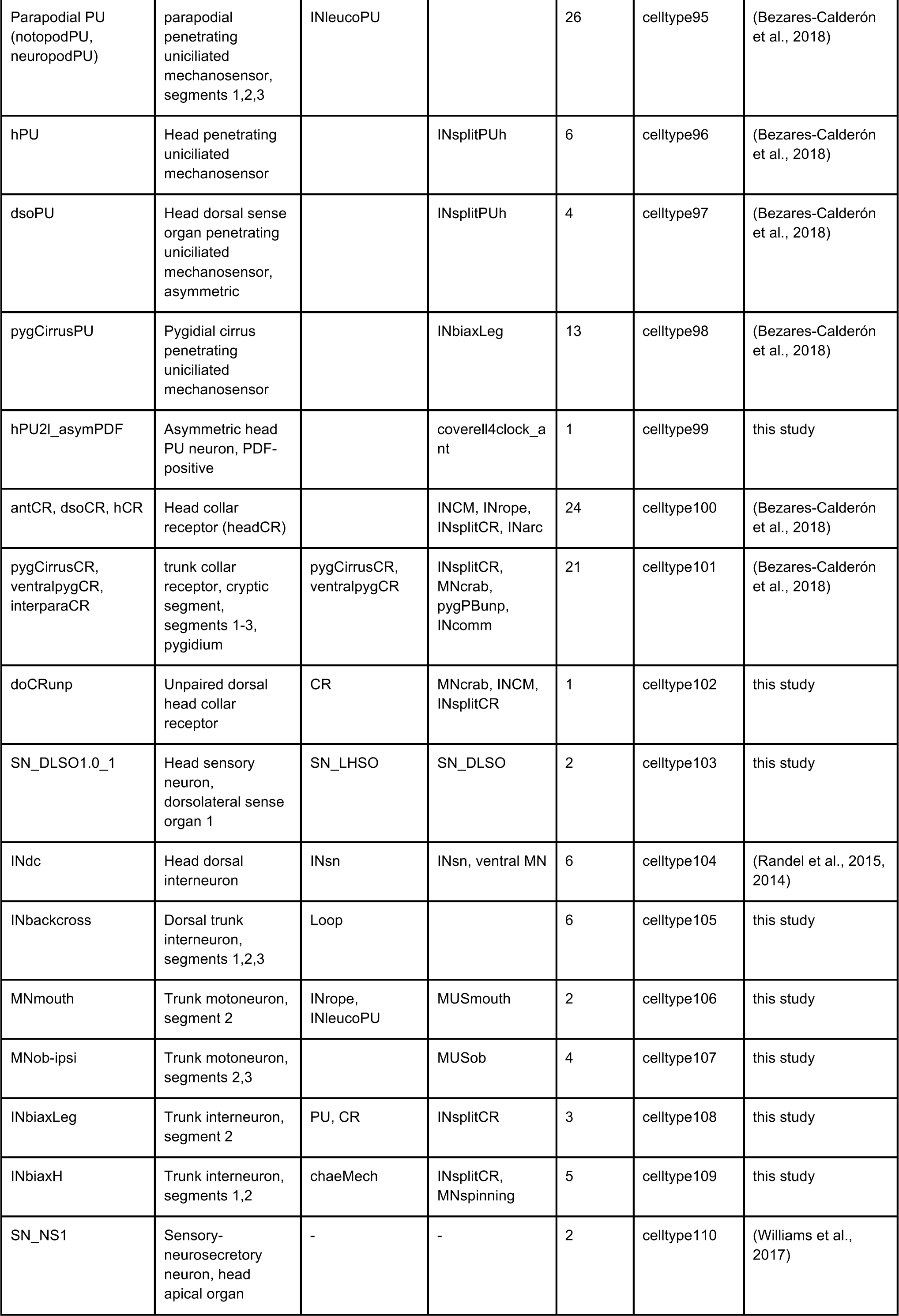
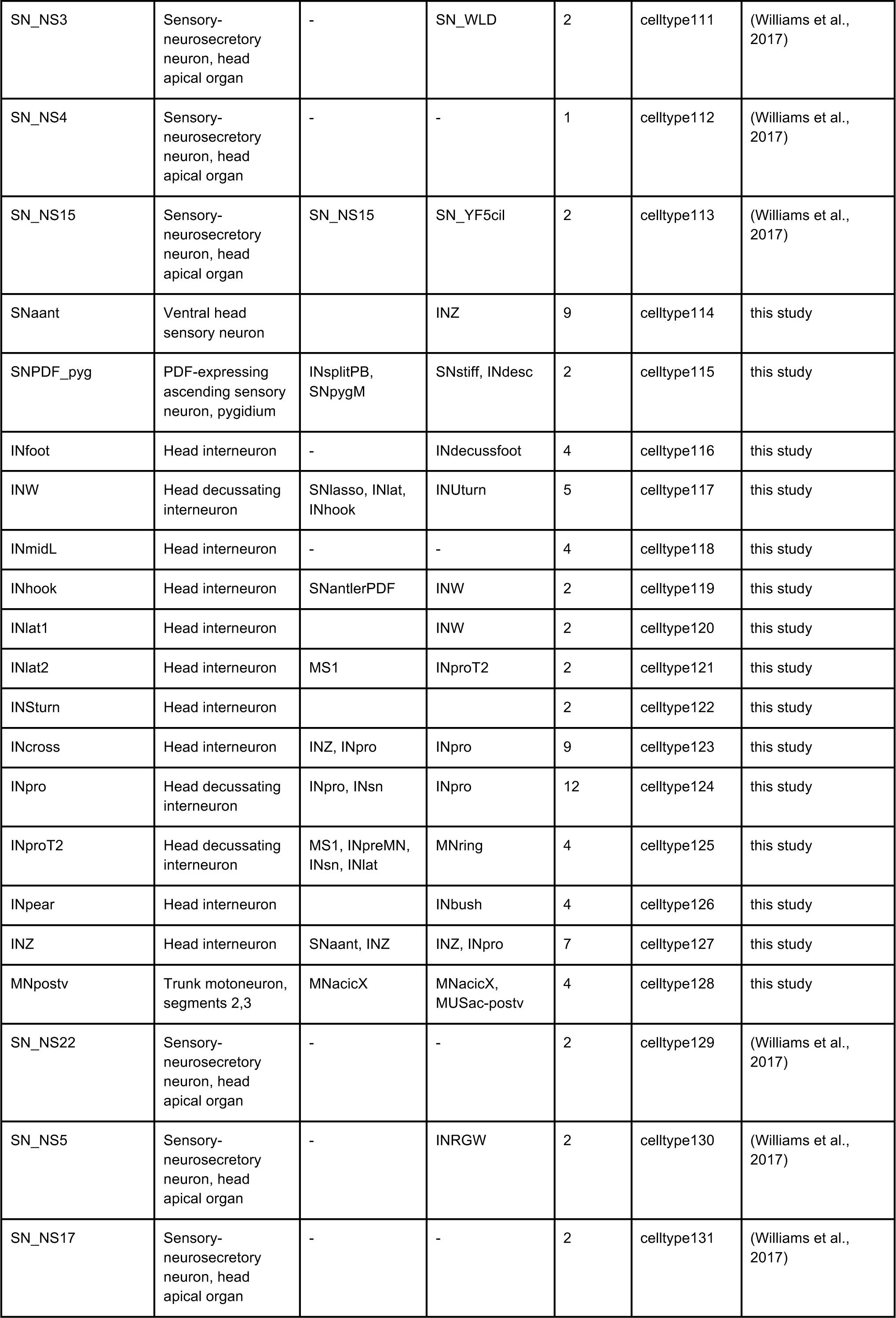
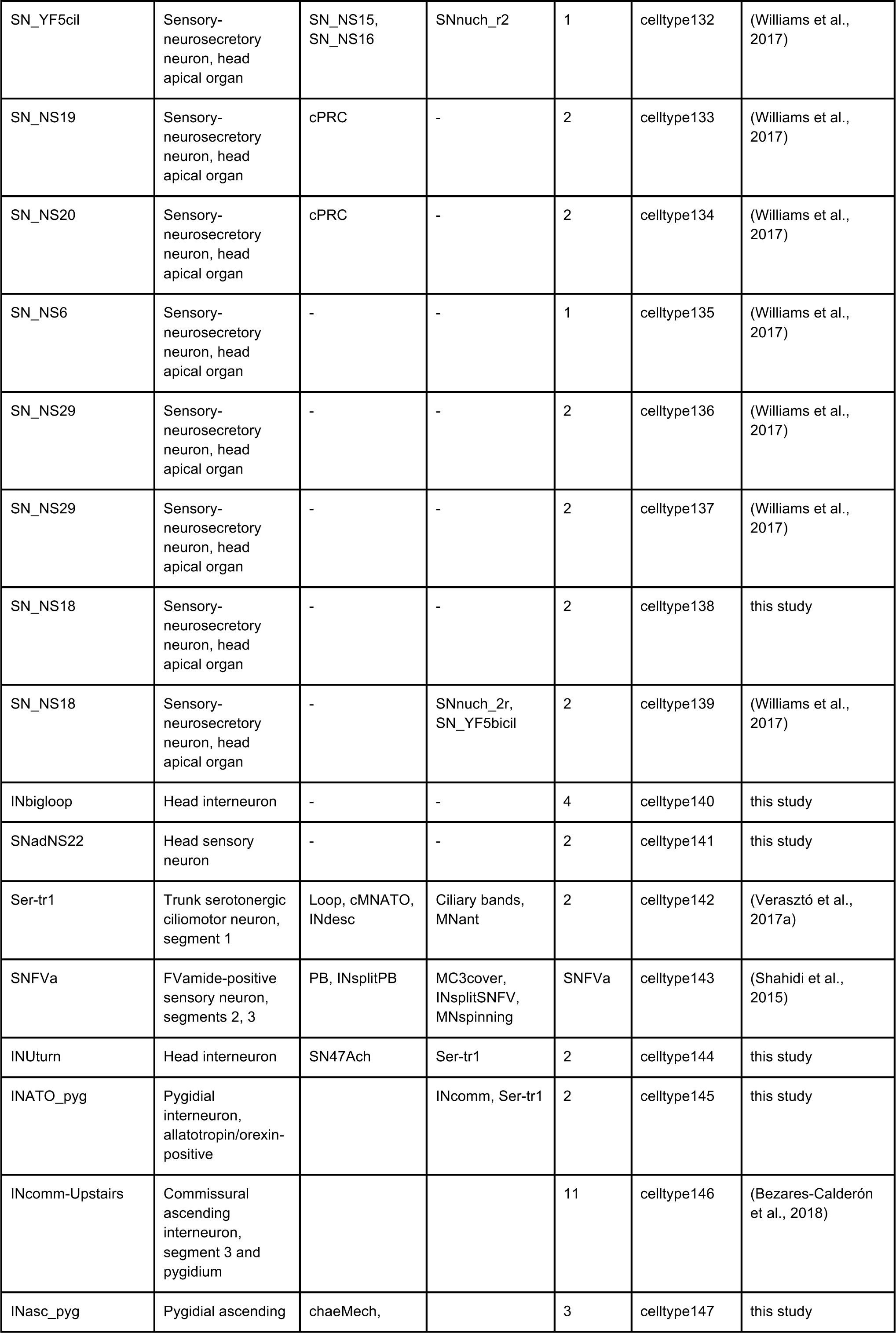
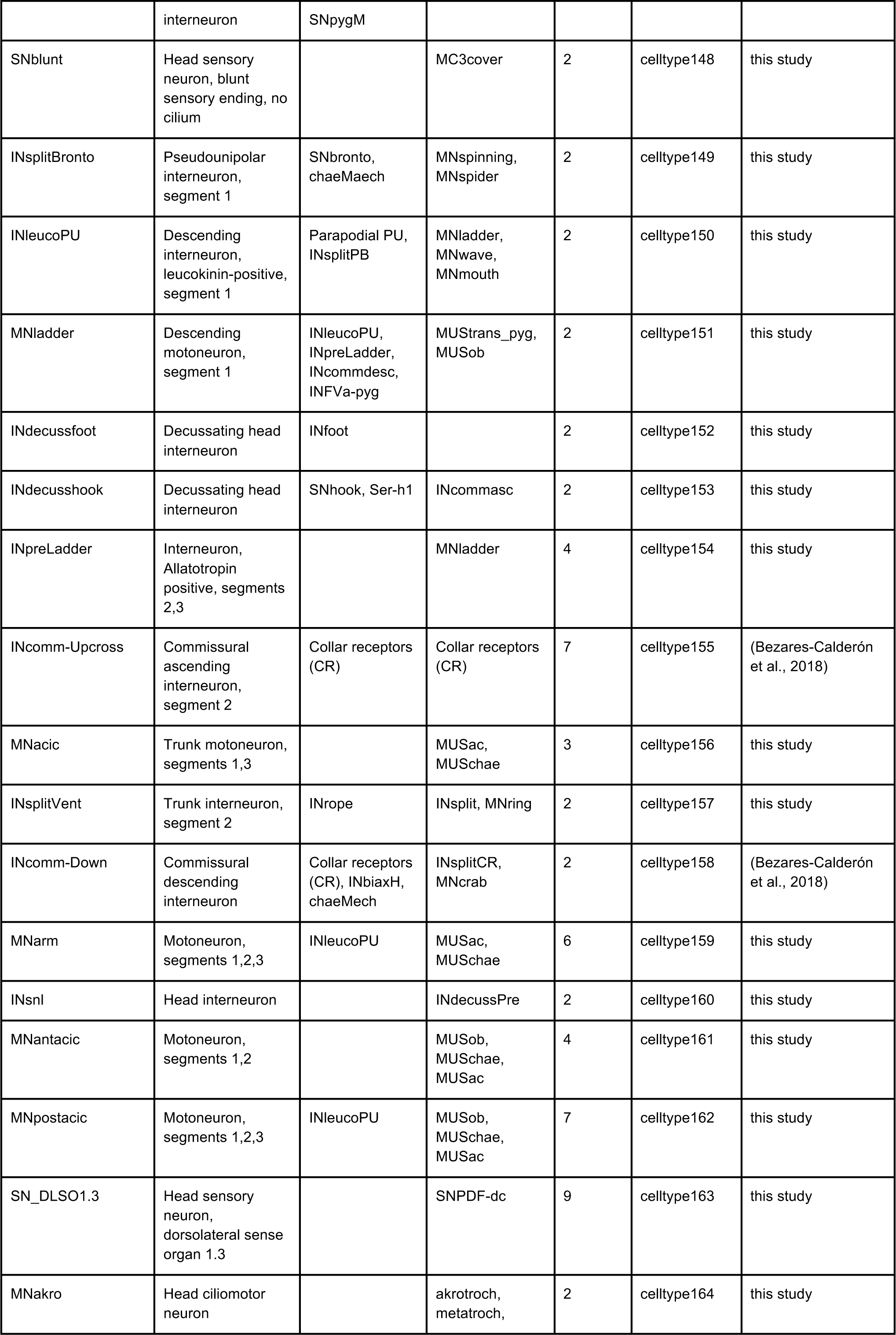
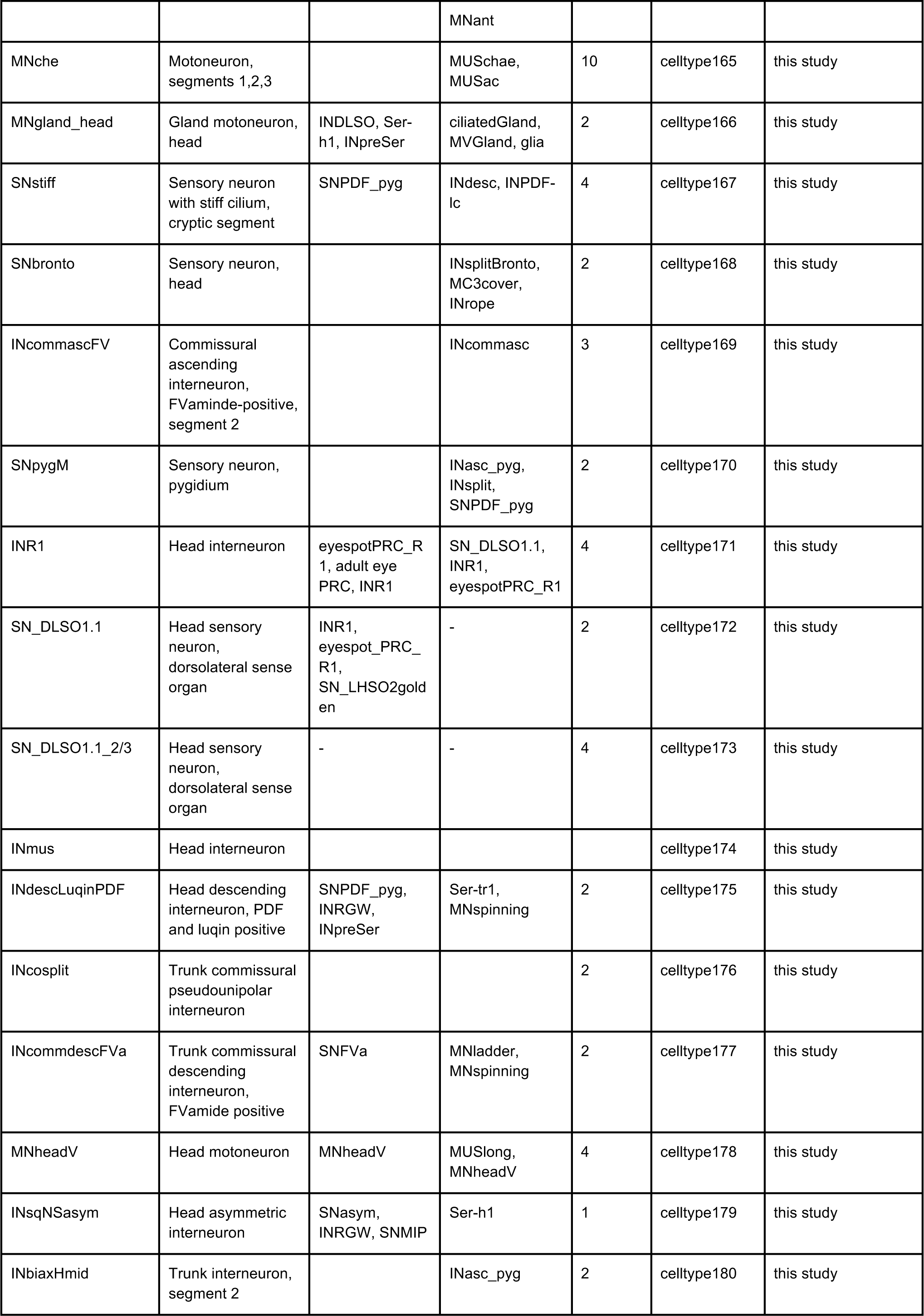
Neuronal cell types in the *Platynereis* three-day-old larval connectome. Cell types are listed in an order matching their CATMAID annotation (celltype1-180).

### Neuronal morphological cell groups

**Table 2.** Neuronal morphological cell groups in the *Platynereis* three-day-old larva. Cell groups are listed in an order matching their CATMAID annotation (cellgroup1-18).

| Common name | description | Included cell types | # of cells | CATMAID annotation |
| --- | --- | --- | --- | --- |
| INipsiasc | ipsilateral, ascending trunk interneurons | INATO_pyg, INasc_pyg, INFV_pyg | 111 | cellgroup1 |
| INipsidesc | ipsilateral, descending trunk interneurons | INleucoPU | 55 | cellgroup2 |
| INcommasc | commissural, ascending trunk interneurons | INcomm-Upstairs, INcomm-Upcross, INcommasc_FV, INpreLadder, INchaeMech | 114 | cellgroup3 |
| INcommdesc | commissural, descending trunk interneurons | INcommdesc_FVa, INcomm-DownL | 37 | cellgroup4 |
| INcomm | commissural trunk interneurons with no ascending or descending neurite | - | 56 | cellgroup5 |
| INcosplit | commissural, pseudounipolar trunk interneurons | INcosplit_sg2 | 24 | cellgroup6 |
| INsplit | ipsilateral, pseudounipolar trunk interneurons | INCM, INsplitCR, INsplitCRATO, INsplitPB, INsplitPBant, INsplitPBant_RF/Ya, INsplitPUh, INsplitVent, INsplitBronto | 114 | cellgroup7 |
| SN_LSO | lateral head sense organ sensory neurons | SN_NS18 | 27 | cellgroup8 |
| SNpyg | pygidial sensory neurons | Pygidial cirrus CR, PB and PU, SNPDF, SNpygM, | 76 | cellgroup9 |
| SNpalp | palp immature sensory neurons | - | 66 | cellgroup10 |
| SN_HSG3 | immature sensory neurons of head sensory group 3 | - | 12 | cellgroup11 |
| INdecuss | decussating head interneurons | INdecussfoot, INdecusshook, INdecussPre, INW, INarc, INCrossbow, INpro | 69 | cellgroup12 |
| INdesc | descending head interneurons | INrope, INbigloop, INMC3, INUturn, INdescLuqinPDF | 33 | cellgroup13 |
| INbush | ventral head interneurons | - | 6 | cellgroup14 |
| SNant, antCR, antPU | antennal sensory neurons, mostly immature | antCR, antPU | 88 | cellgroup15 |
| SNcir | sensory neurons of cirri, mostly immature | - | 60 | cellgroup16 |
| SNstomod | stomodaeal sensory neurons, immature | - | 51 | cellgroup17 |
| SNimmature_sg0 | cryptic segment dorsal sensory organ, immature | - | 15 | cellgroup18 |

### Non-neuronal cell types

**Table 3.**
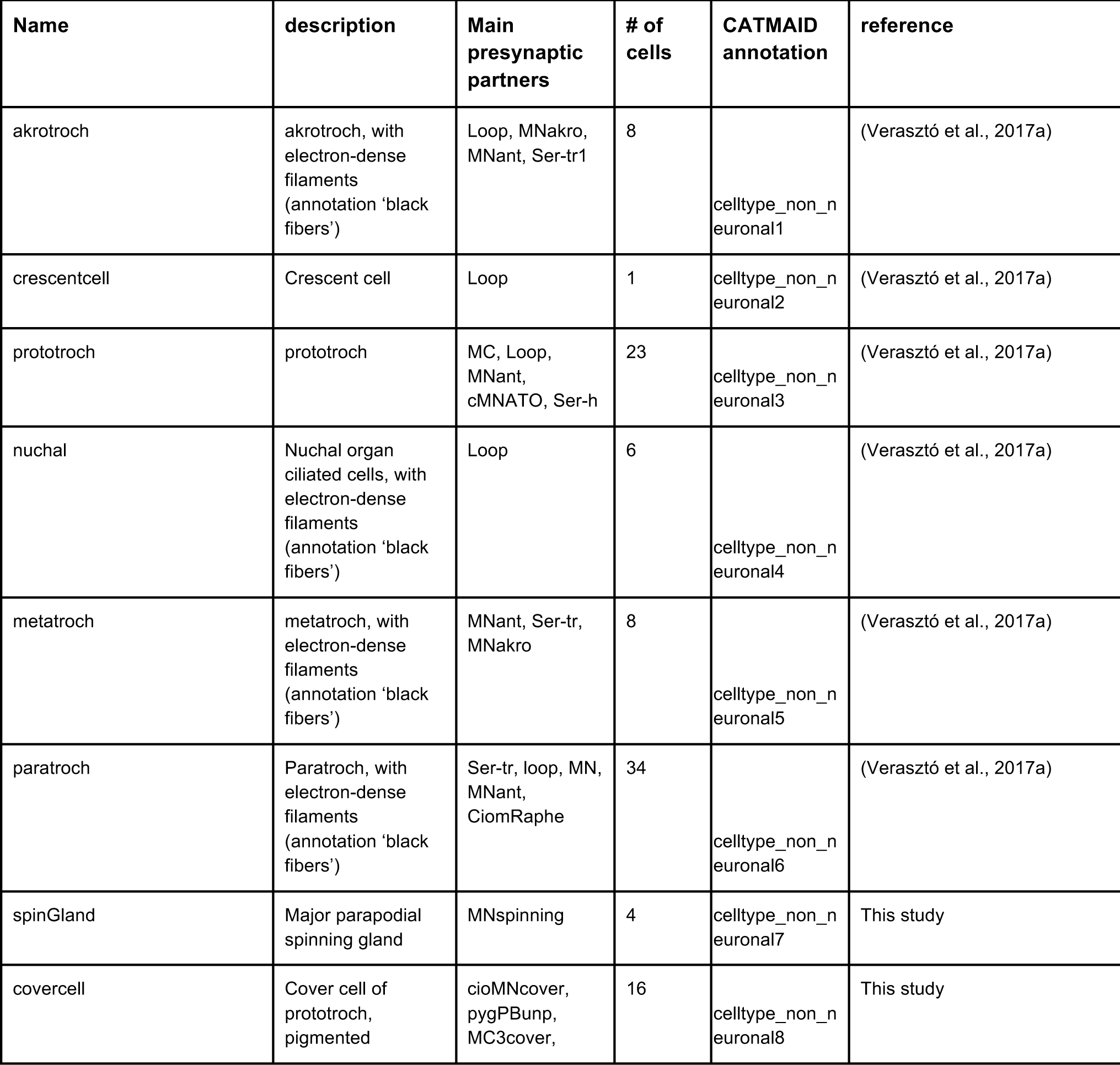
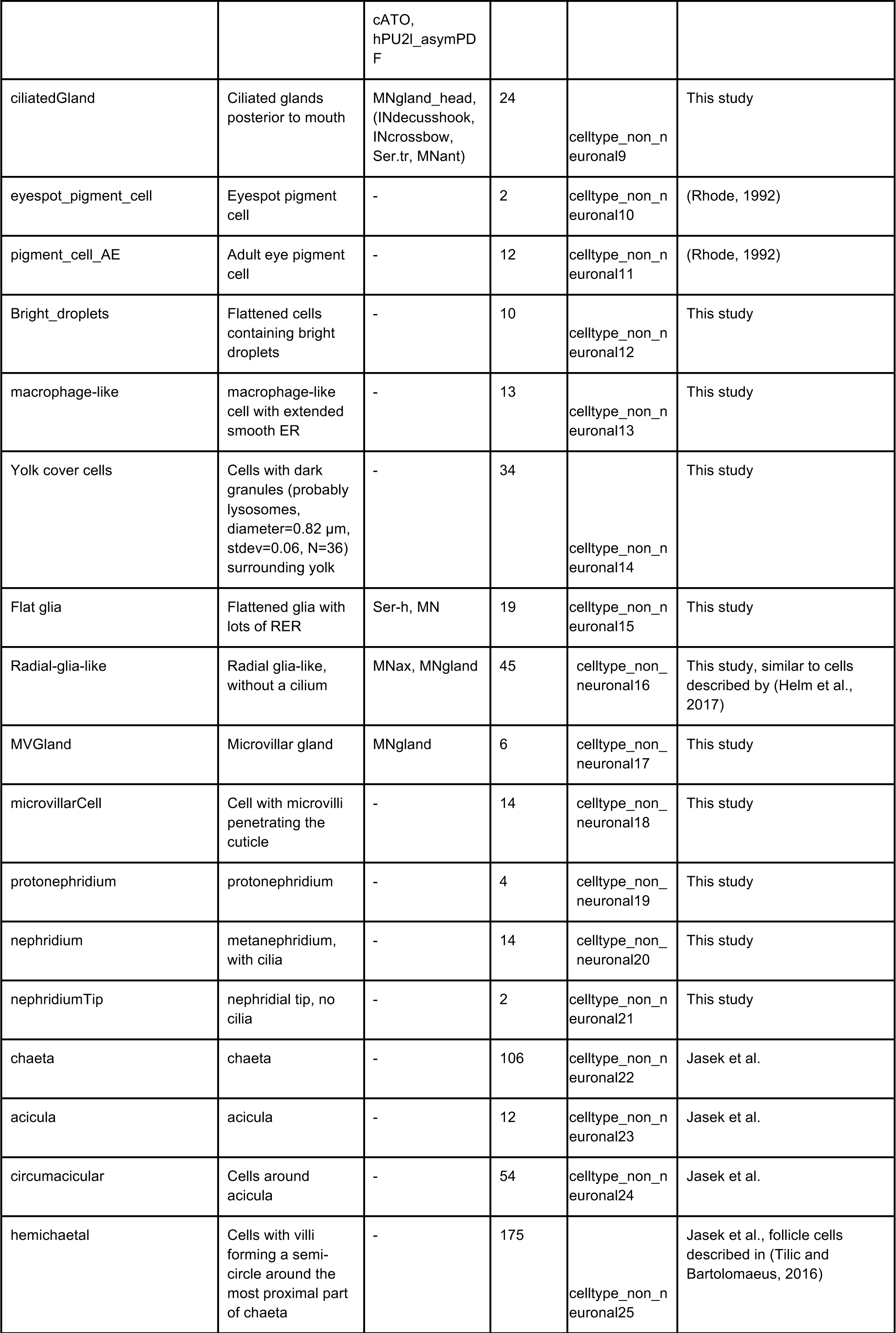
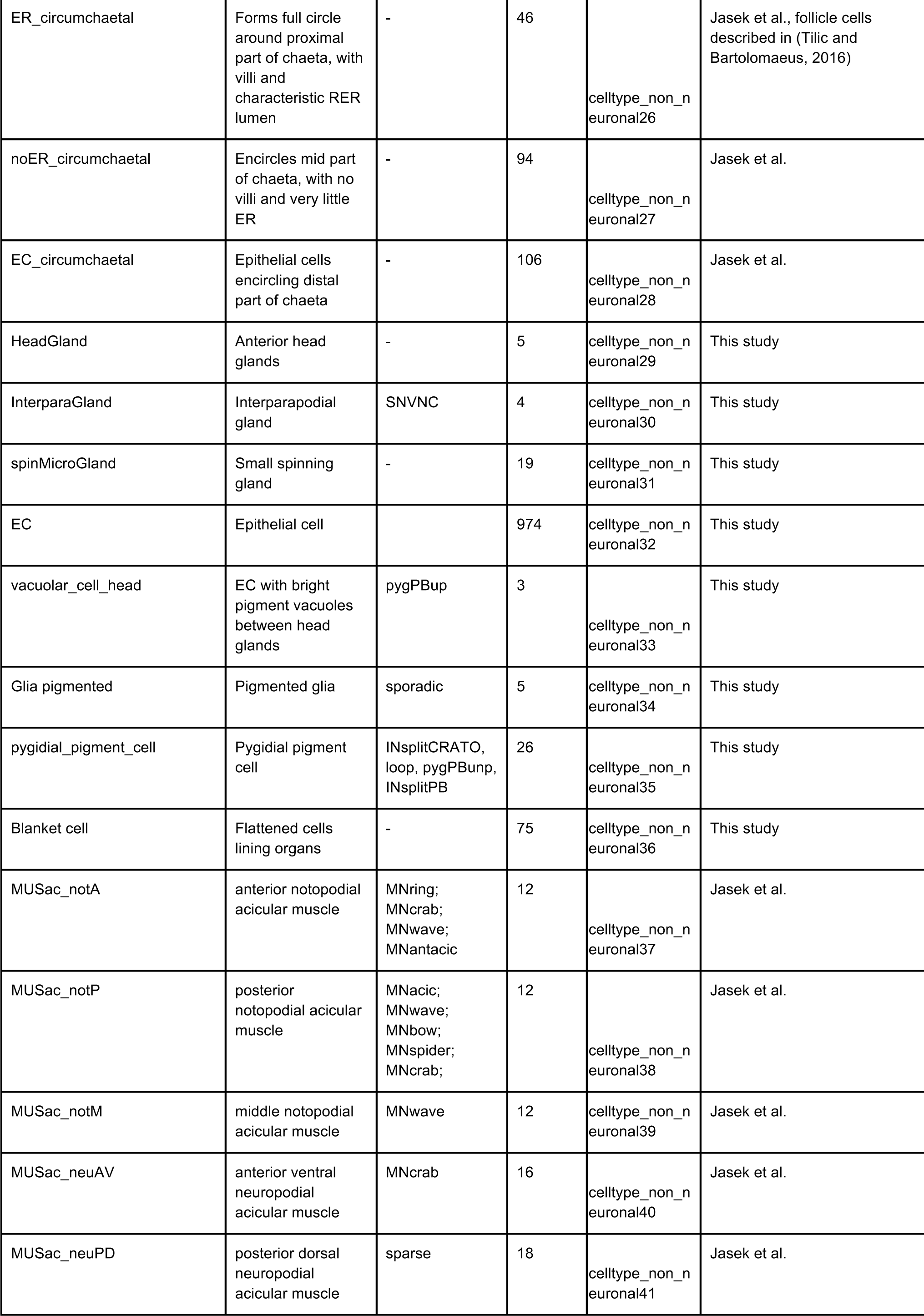
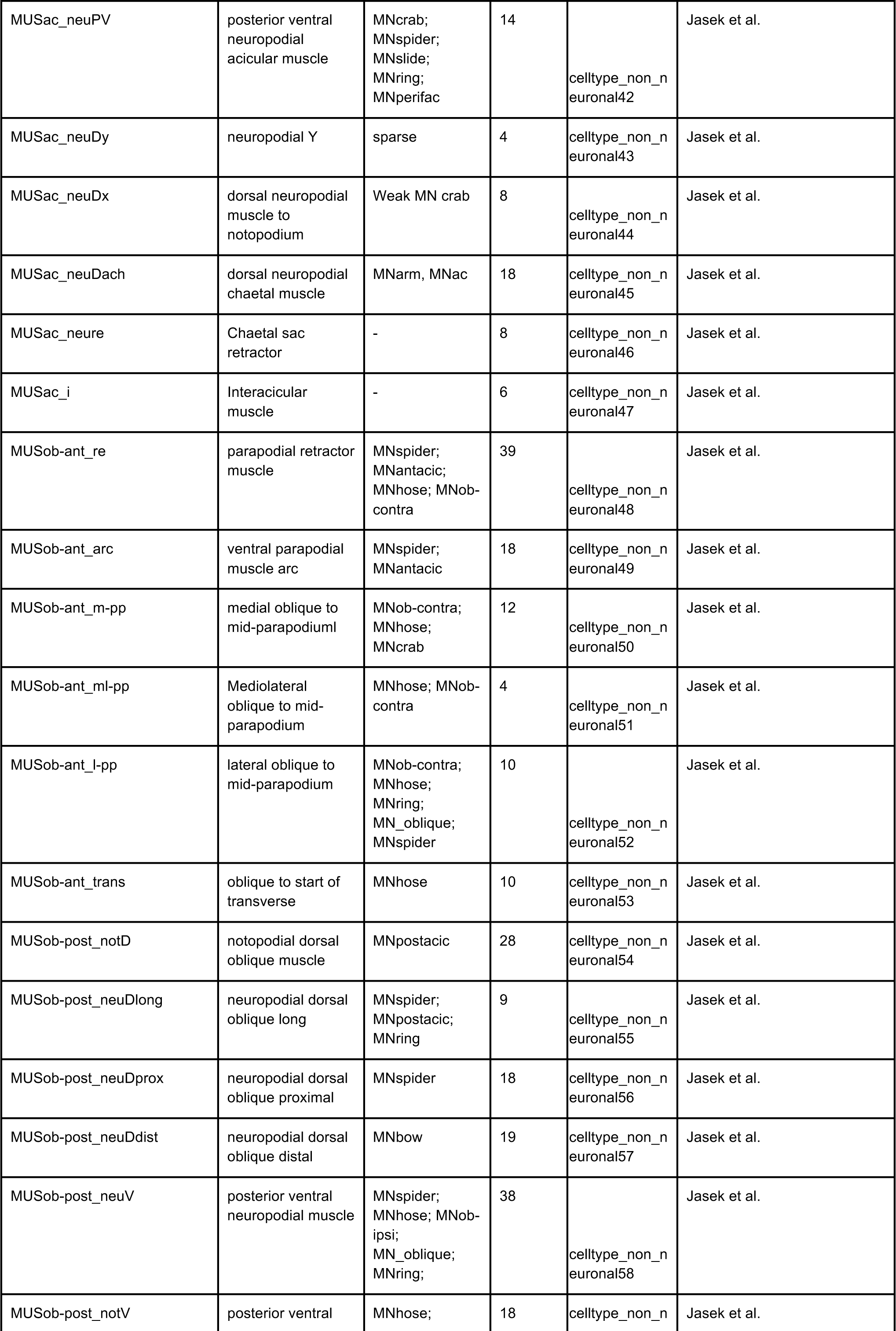
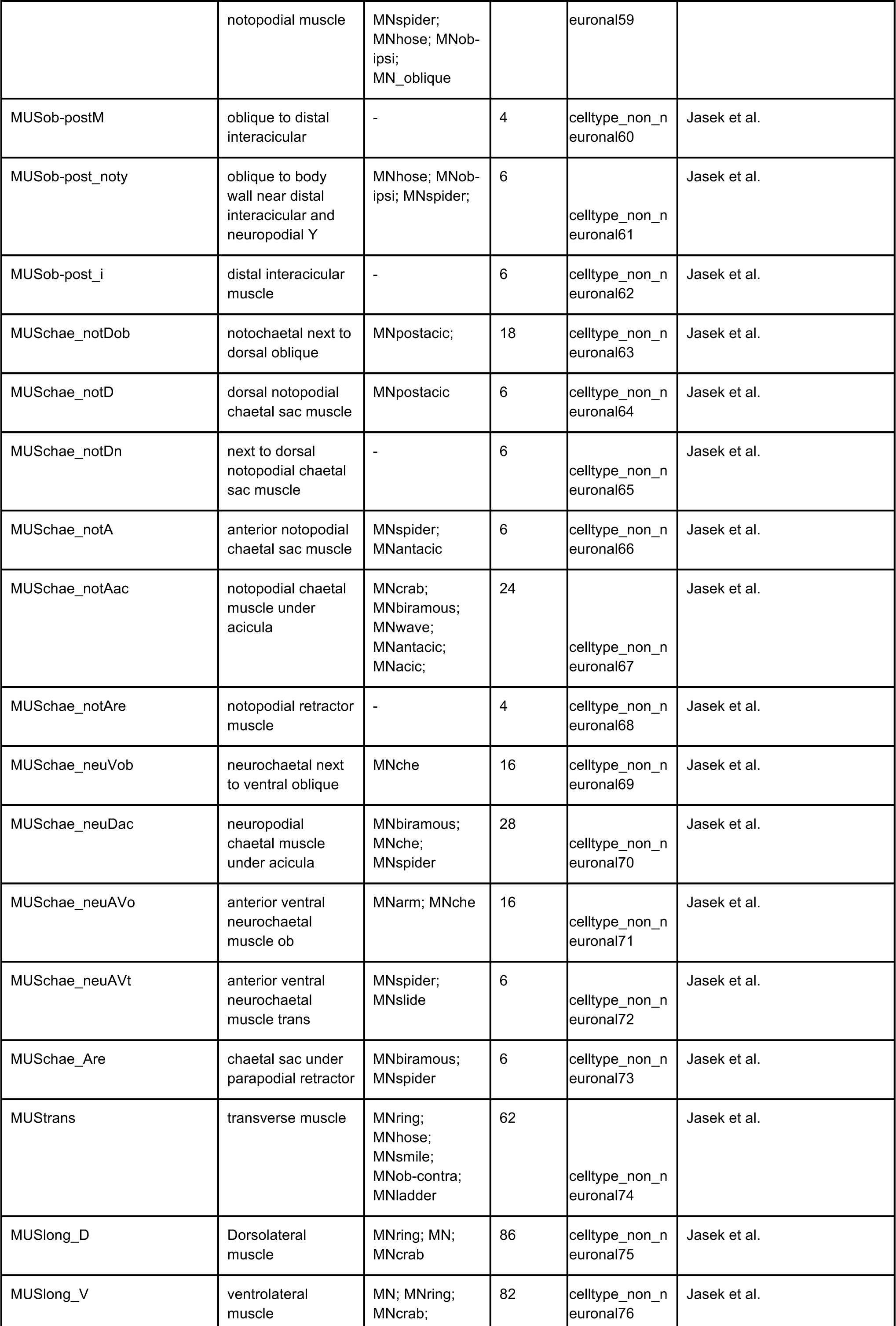
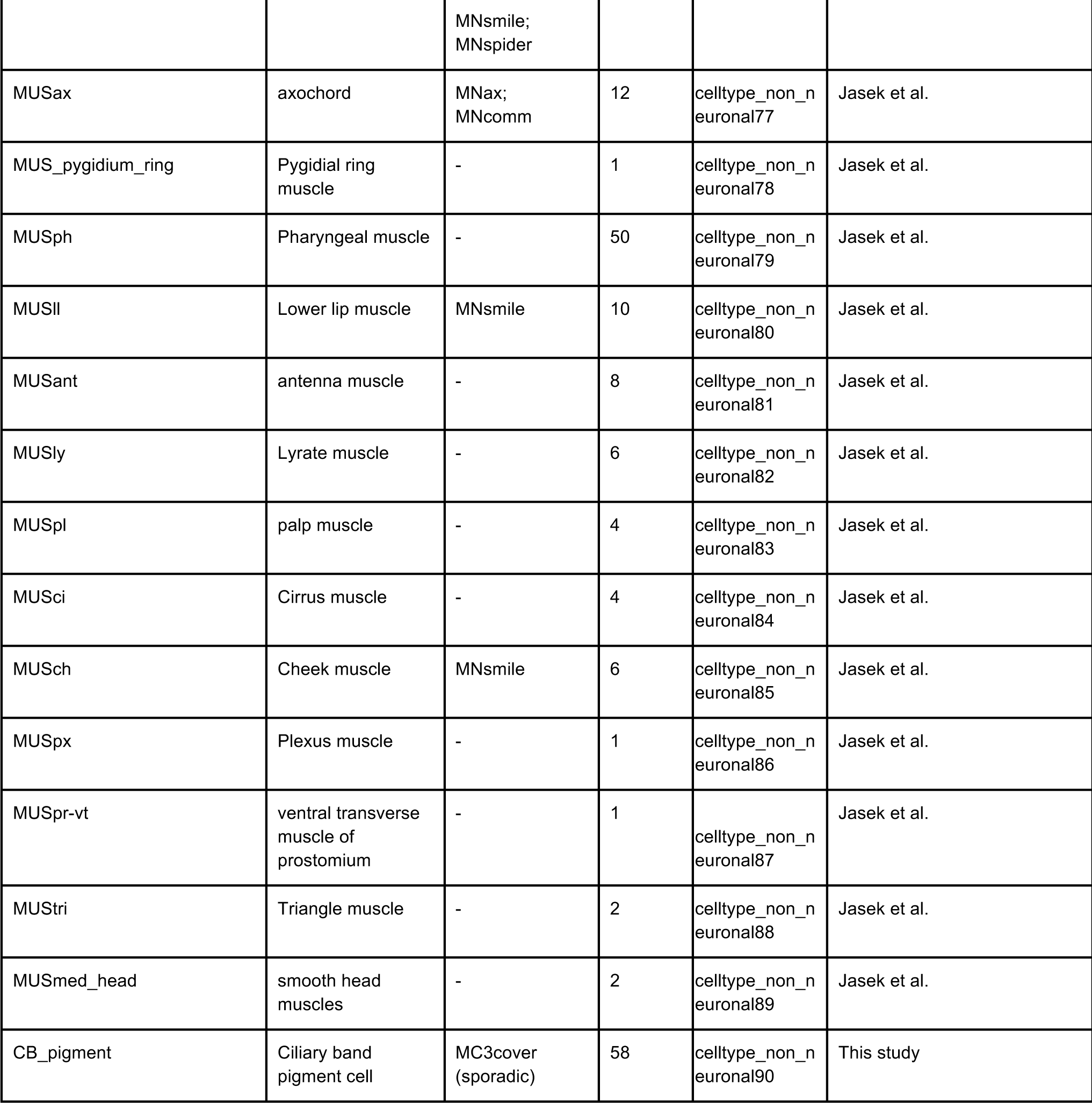
Non-neuronal cell types in the *Platynereis* three-day-old larva. Cell types are listed in an order matching their CATMAID annotation (celltype_non_neuronal1-89).

**Table 4.** CATMAID accessory project layers related to the main HT9-4 (NAOMI) stack. Summary of the extra project stacks that were created by re-imaging the main data set at higher resolutions. This was needed to resolve some inconsistencies and hard-to-trace layers.

| CATMAID projects | Number of<br>layers<br>imaged (z) | Layer dimensions (x,y)<br>in pixels | Resolution (x,y,z) nm/pixels |
| --- | --- | --- | --- |
| HT-4_Naomi_project | 4845 | 25792, 28800 | 5.7, 5.7, 40 |
| Plexus_HT-4_Naomi_project__372-4013 | 1407 | 43920, 40968 | 2.2, 2.2, 40 |
| Immuno_HT-4_Naomi_project | 13 | 50000, 52000 | 2.2, 2.2, 40 |
| Jump_864_HT-4_Naomi_project__853-873 | 21 | 60000, 60000 | 2.2, 2.2, 40 |
| Jump_2800_HT-4_Naomi_project__2762-2818 | 57 | 44702, 26371 | 2.2, 2.2, 40 |
| Jump_3725_HT-4_Naomi_project__3719-3728 | 10 | 24000, 26000 | 3.7, 3.7, 40 |

---

**Figure 1 – figure supplement 1.**
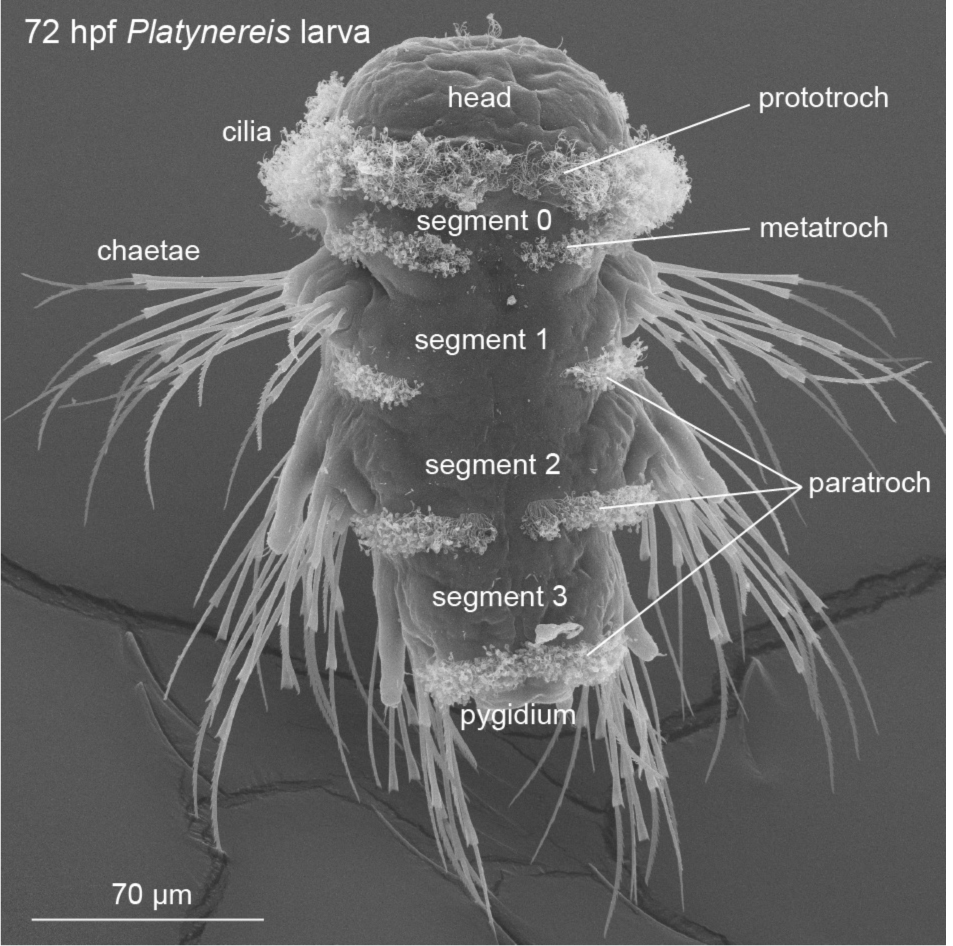
Anatomy of the three-day-old *Platynereis* larva. Scanning EM image of a three-day-old *Platynereis* larva (72 hours post fertilisation) with the main body regions labelled.

**Figure 1 – figure supplement 2.**
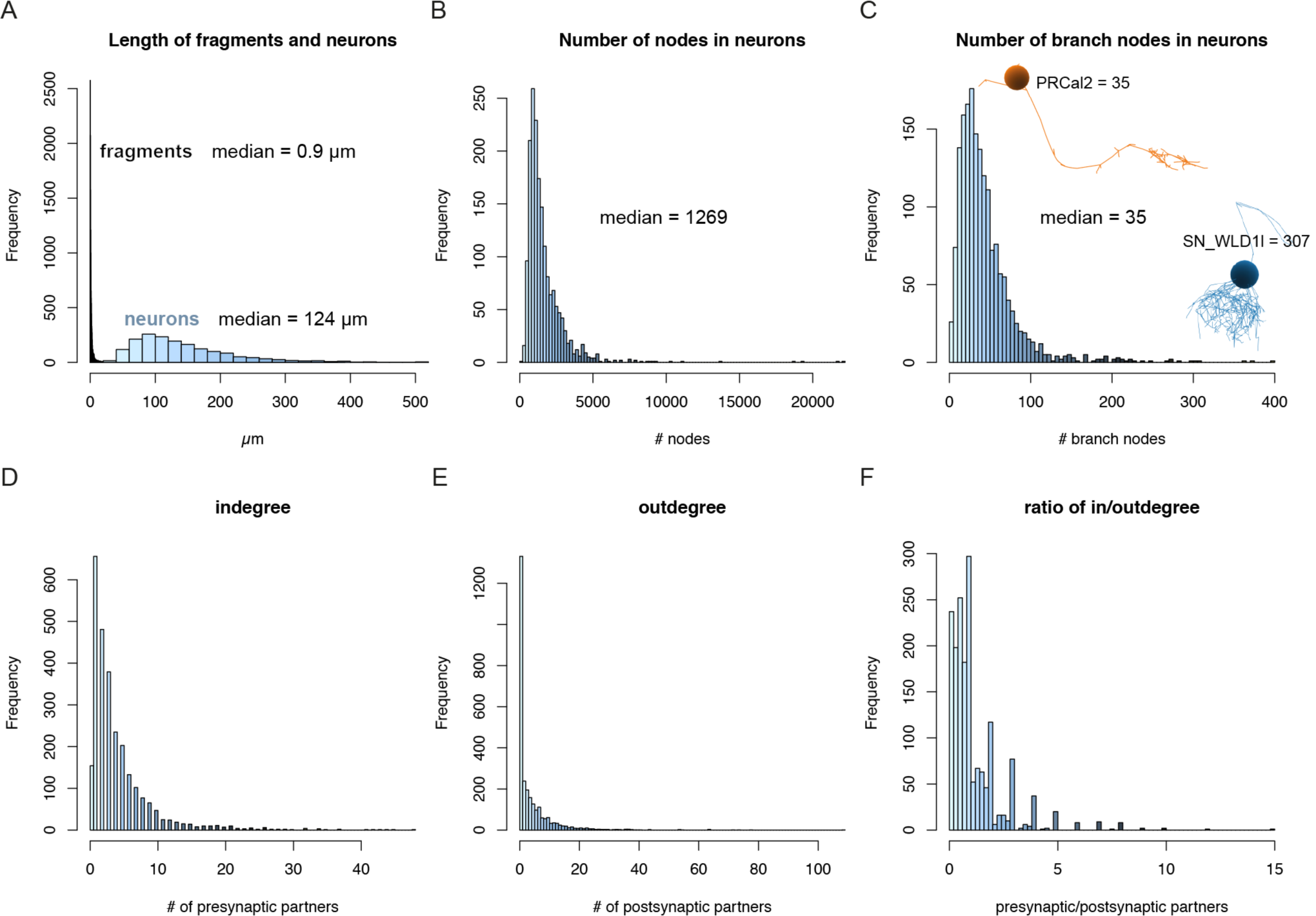
Skeleton statistics of the *Platynereis* larval connectome. (A) Length distribution of fragments and neurons with soma in the connectome set. (B) Histogram of the number of nodes in connectome neurons. (C) Histogram of the number of branch nodes in the connectome neurons. The images show example neurons with a simple (PRCal2) and a highly branched skeleton (SN_WLD1l). (D) Indigree distribution of all cells in the connectome set. (E) Outdegree distribution of all cells in the connectome set. (F) The distribution of the ratio of indegree to outdegree for all cells in the connectome set.

**Figure 1 – figure supplement 3.**
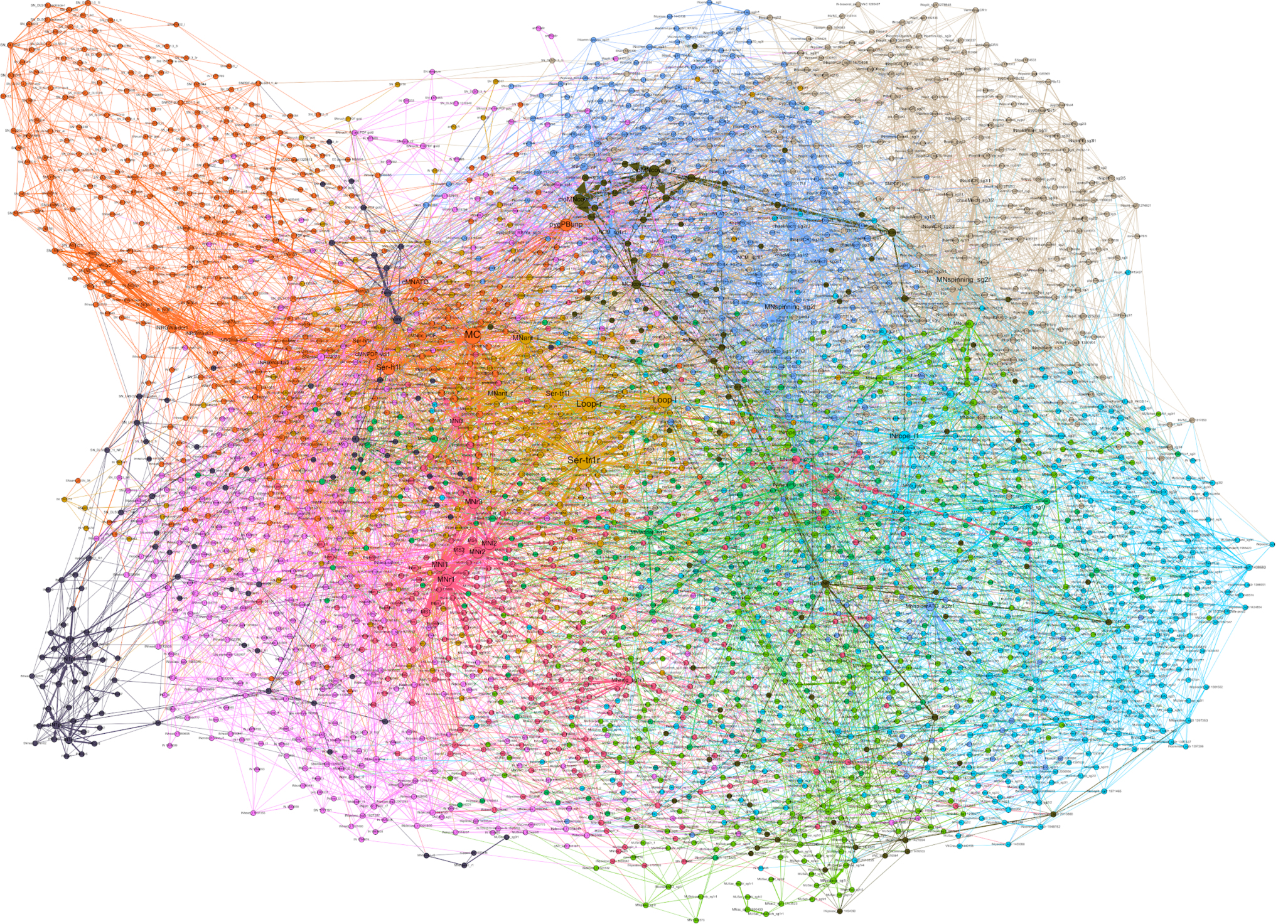
The *Platynereis* larval connectome with node labels. Graph representation of the connectome with nodes representing single cells and edges synaptic connectivity. The graph includes 2,728 nodes and 11,403 edges.

**Figure 3 – figure supplement 1.**
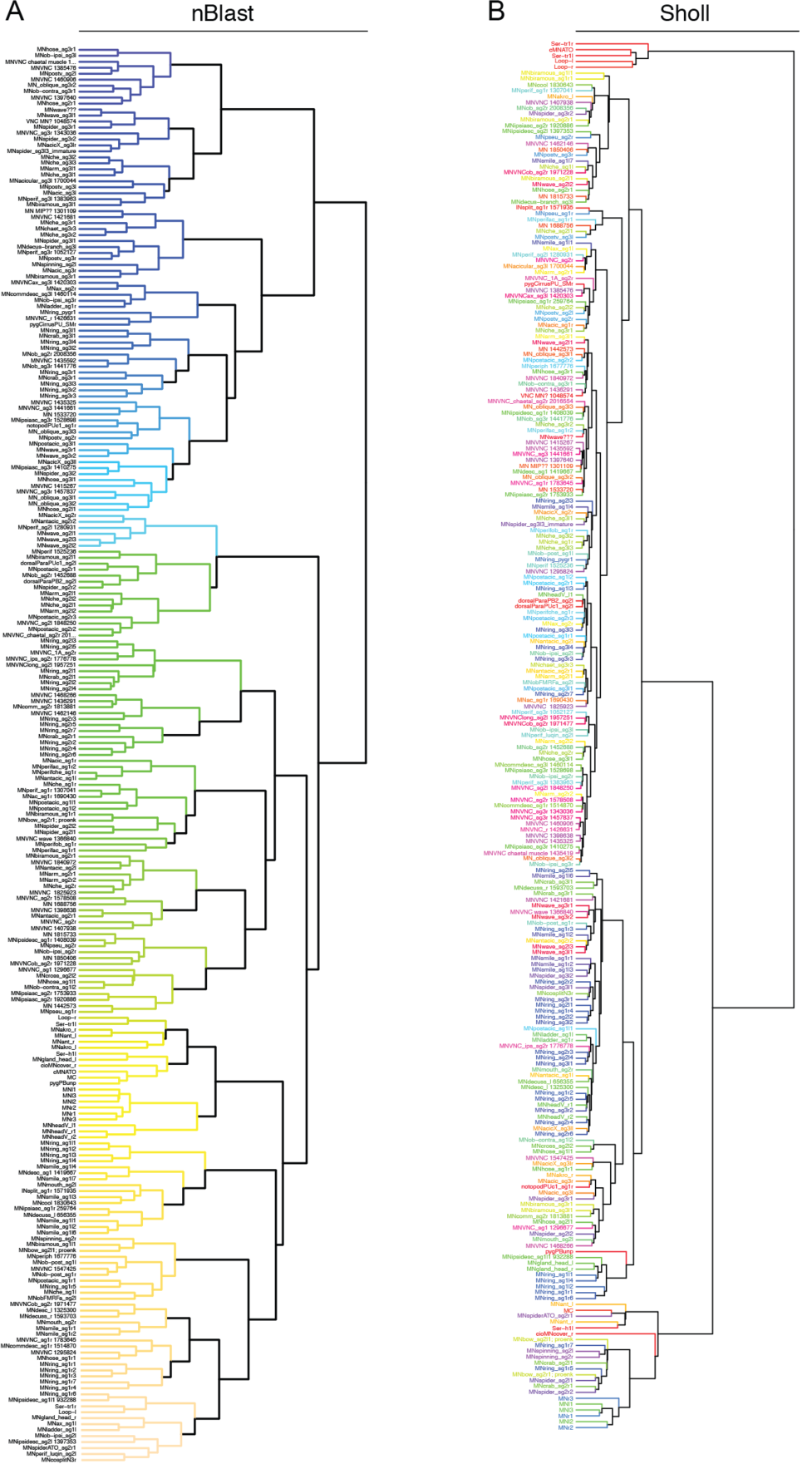
Clustering of motoneurons based on NBlast and Sholl analysis. (A) Dendrogram of 237 motoneurons based on NBLAST scores. (B) Dendrogram of 237 motoneurons based on Sholl analysis scores. In B neurons that we defined as cell types have similar colours.

**Figure 3 – figure supplement 2.**
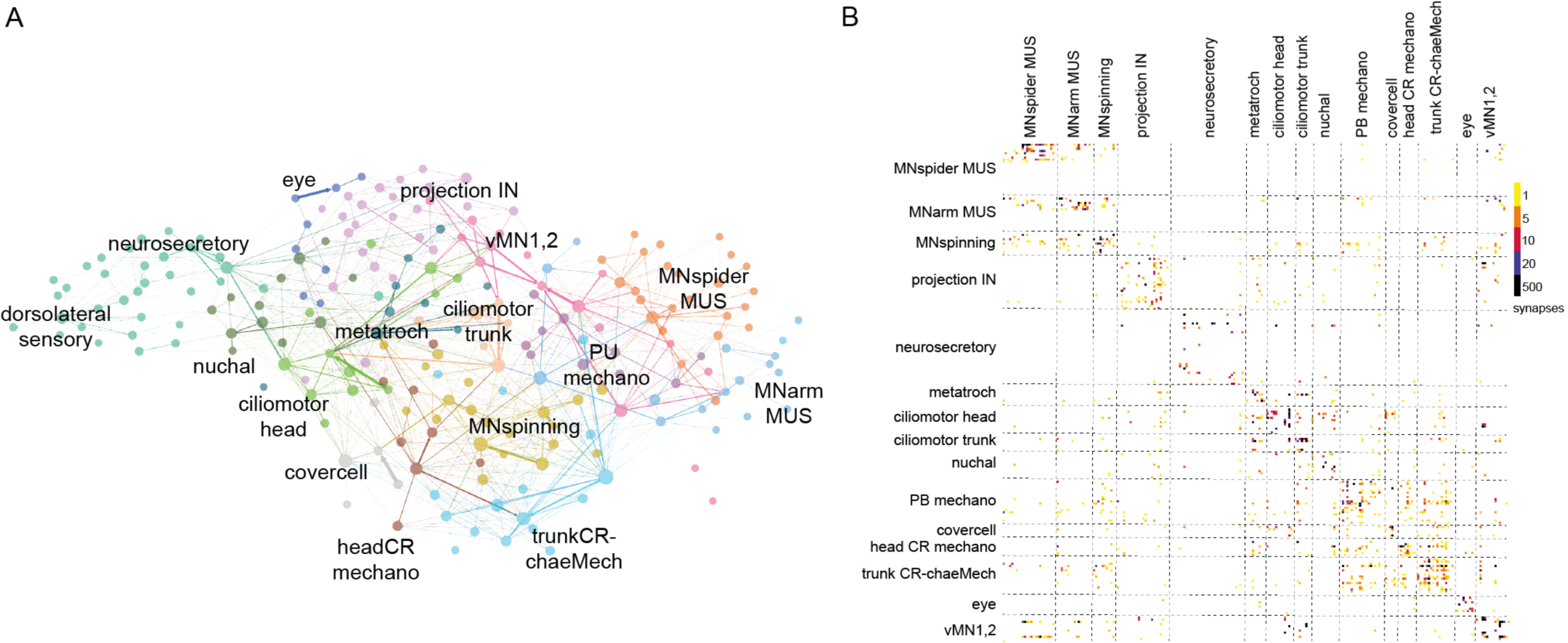
Modules in the cell-type connectome. (A) Modules in the cell-type connectome. (B) Connectivity matrix of the graph in Figure 3, dashed lines delineate the modules coloured in A.

**Figure 3 – figure supplement 3.**
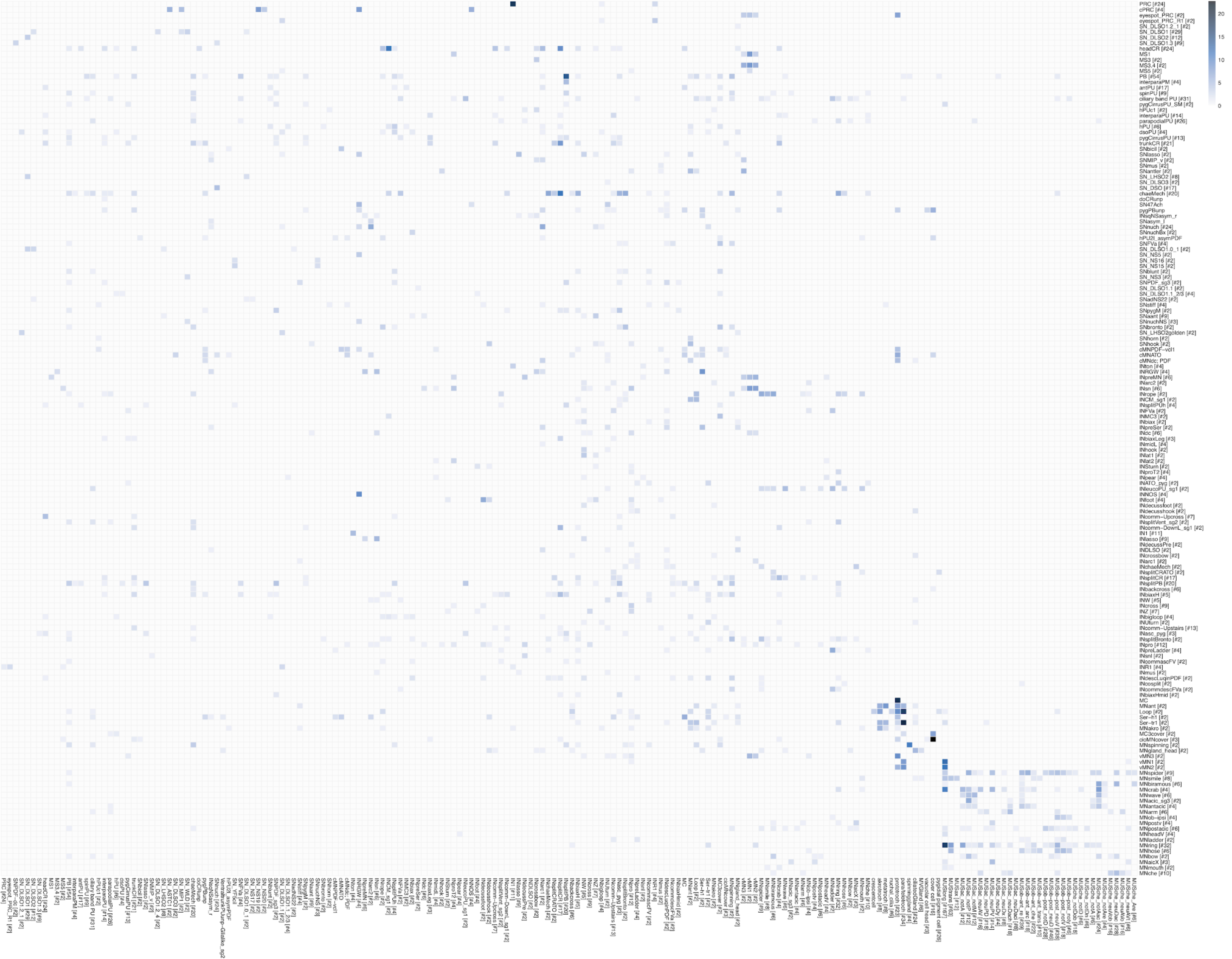
Connectivity matrix of the cell-type connectome. Grouped connectivity matrix of cell types. The number of cells of the same type in a group is shown in square brackets. The scale represents the square root of the number of synapses. Presynaptic groups are shown on the right side.

**Figure 3 – figure supplement 4.**
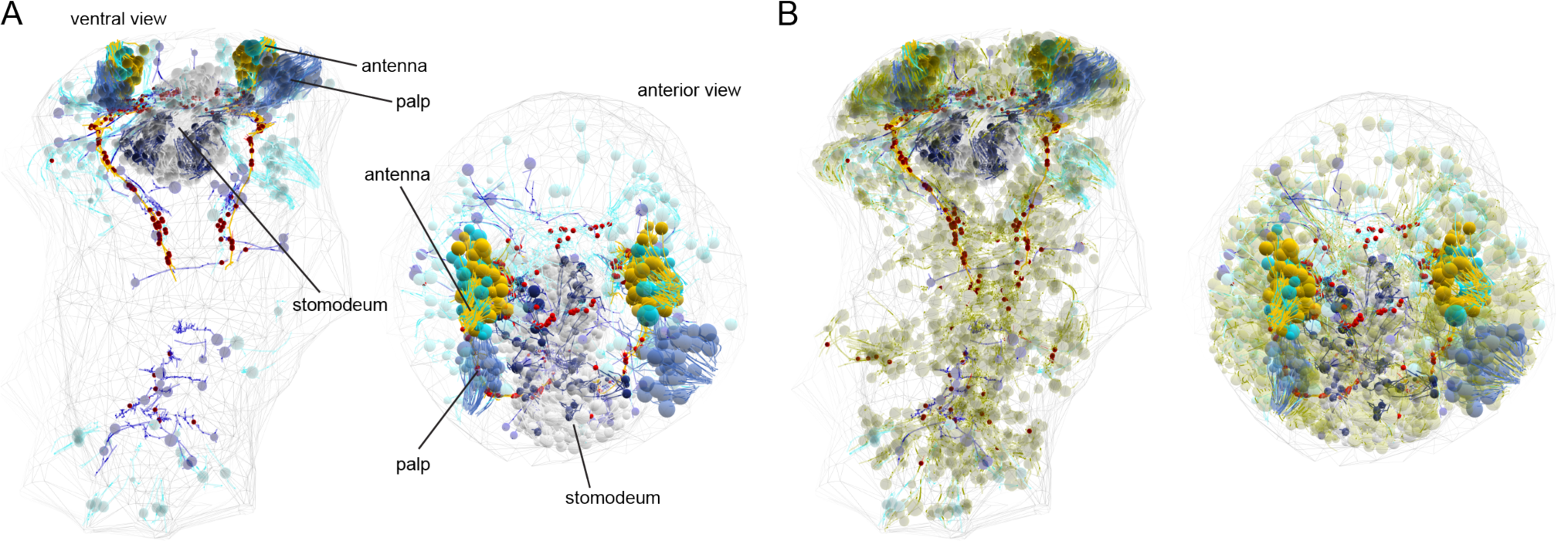
Developing sensory neurons. (A) Reconstruction of antennal, palp and stomodeal cells. The cyan cells in the antennal group are differentiated. All other cells are immature, with immature sensory dendrites and few synapses. Ventral and anterior views. Presynaptic sites are shown as red dots. (B) The same cells shown together with 1,692 non-differentiated cells that putatively belong to the neuronal lineage.

**Figure 7 – figure supplement 1.**
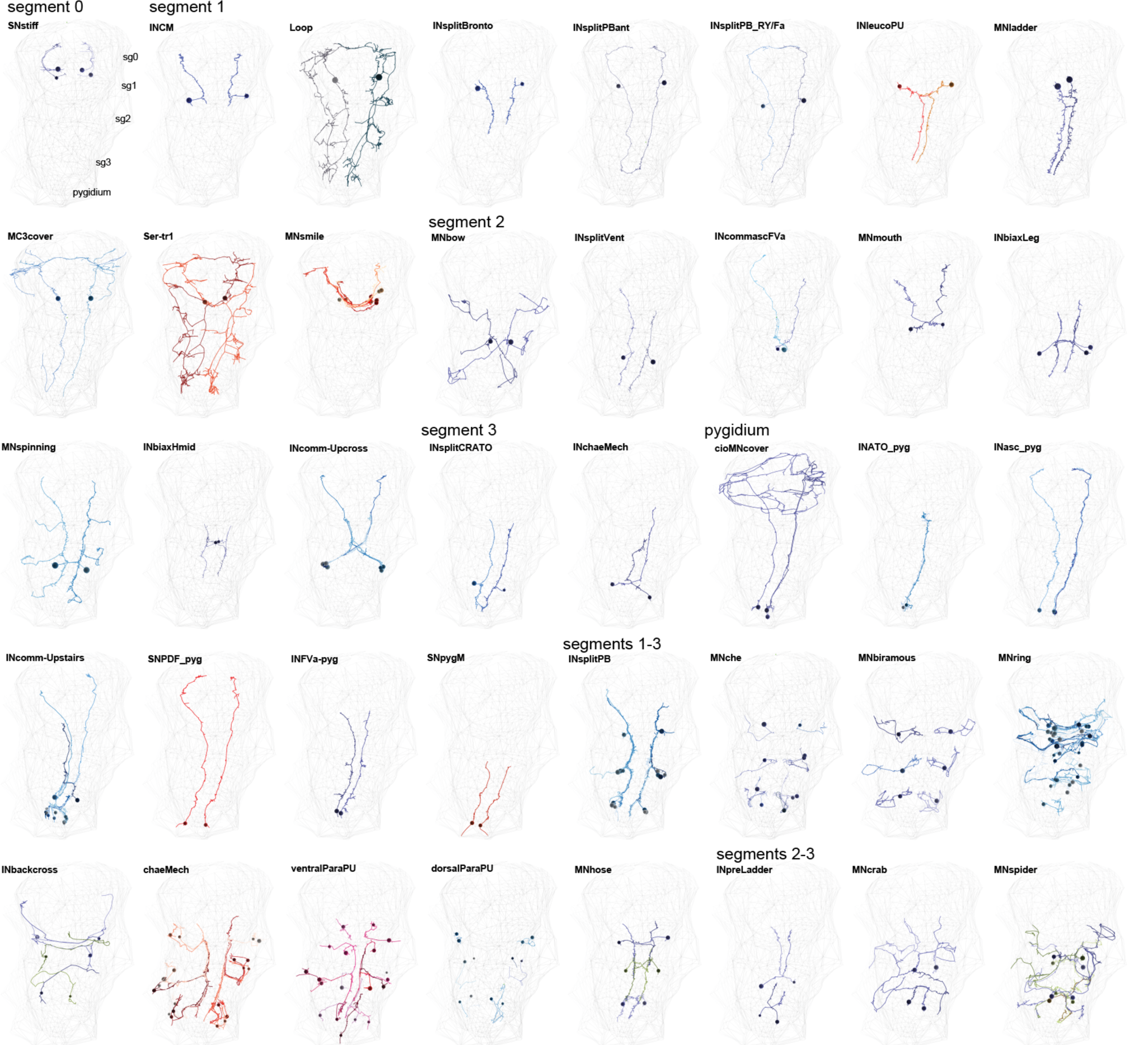
Segment-specific and segmentally iterated cell types. EM reconstructions of sensory, inter- and motor neuron types present in different combinations of trunk segments.

**Figure 8 – figure supplement 1.**
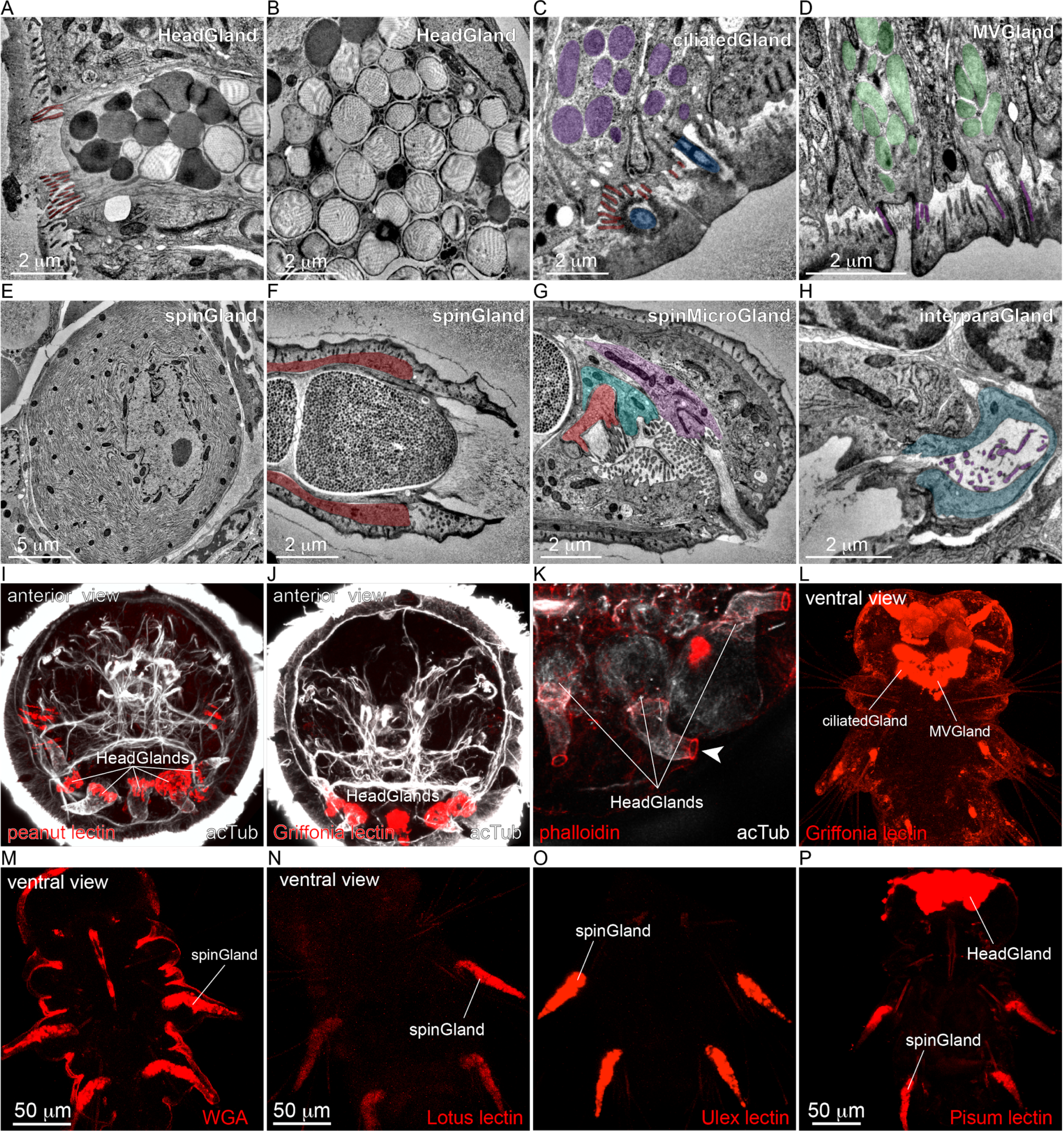
Ultrastructure and lectin reactivity of exocrine glands. (A) TEM image of the secretory pore of a head gland cell. (B) Secretory vesicles in a head gland cell. (C) TEM image of a ciliated gland cell. The cilium, microvilli and vesicles are highlighted. (D) A microvillar gland cell. The microvilli and vesicles are highlighted. (E) Soma of a spinning gland cell with the extended ER. (F) Secretory pore of a spinning gland cell. (G) Secretory pore of three spin micro gland cells adjacent to the pore of the spinning gland cell. (H) Secretory pore of an interparapodial gland cell. (I) Staining of headGland cells with peanut lectin in a two-day-old larva, anterior view. (J) Staining of headGland cells with Griffonia lectin in a two-day-old larva, anterior view. (K) Phalloidin staining of the secretory pore of head gland cells. (L) Staining of ciliateGland and MVGland cells with Griffonia lectin in a three-day-old larva, ventral view. (M) Staining of spinGland cells with wheat germ agglutinin in a three-day-old larva, ventral view. (N) Staining of spinGland cells with Lotus lectin in a three-day-old larva, ventral view. (O) Staining of spinGland cells with Ulex lectin in a three-day-old larva, ventral view. (P) Staining of spinGland and headGland cells with Pisum lectin in a three-day-old larva, ventral view.

**Figure 8 – figure supplement 2.**
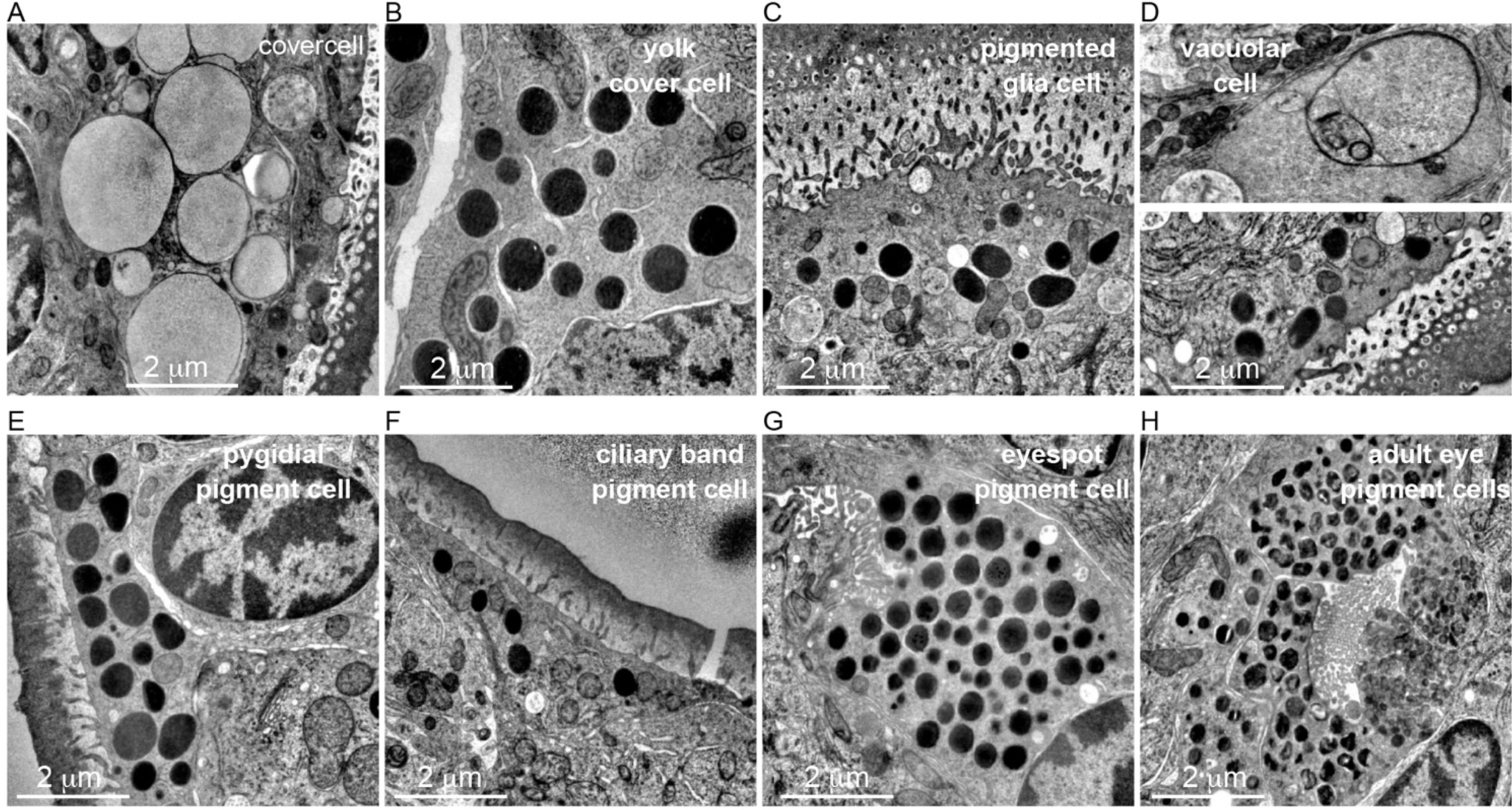
Ultrastructure of pigment cells. (A) TEM image of a covercell with the large pigment vacuoles. (B) TEM image of a yolk cover cell with large granules. (C) TEM image of a pigmented glia cell. (D) TEM images of a vacuolar cell. (E) TEM image of a pygidial pigment cell. (F) TEM image of a ciliary band pigment cell. (G) TEM image of a pigment cell from the eyespot. (H) TEM image of a pigment cell from the adult eye.

**Figure 10 – figure supplement 1.**
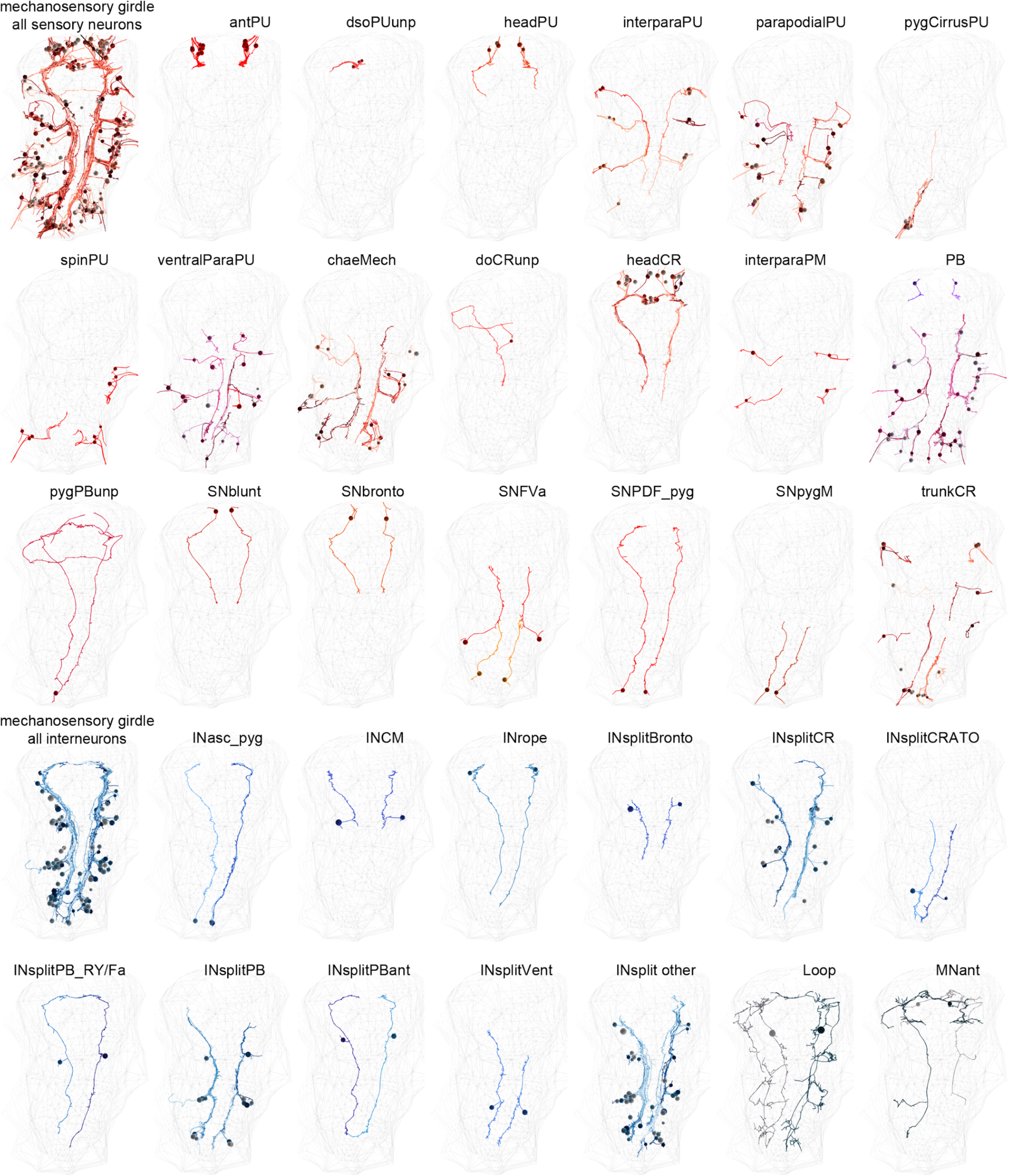
Cell types in the mechanosensory girdle. EM reconstructions of sensory, inter- and motor neuron types that form the mechanosensory girdle.

